# The transition from spontaneous to preparatory movement in infancy

**DOI:** 10.64898/2025.12.03.691254

**Authors:** Ryo Fujihira, Yuta Shinya, Hama Watanabe, Gentaro Taga

**Author notes:** Corresponding author., (RF), (GT). **Competing interests:** Authors declare that they have no competing interests.

## Abstract

Intention lies at the core of human behavior, allowing flexible and goal-directed interaction with the environment. However, how it emerges in infancy remains poorly understood. We investigated whether 3-month-old infants can initiate intentional actions by simultaneously recording electromyogram, eye gaze, and pupil diameter while they interact with a suspended mobile toy. We found that pupil dilation associated with movement emerged through environmental interaction and gradually shifted to precede movement onset. This shift implies that infants begin to anticipate the timing of their forthcoming movements. Furthermore, this antecedent pupil dilation occurred only with efficient movements characterized by reduced muscle co-contractions. These findings suggest that infants learn to predict outcomes of their own actions and initiate particular movements at self-determined timings, marking the emergence of intentional action.

## Main Text

Intentional action constitutes the foundation of our daily lives. It entails the capacity to initiate actions at self-determined timings while anticipating their consequences on the external world (*1–3*). Although this capacity is considered fundamental for human nature, how intentional action emerges during infancy remains largely unknown. In adults, intentional actions are preceded by a neural activity known as the readiness potential (*4*), which reflects the prediction for action outcomes (*5, 6*). The emergence of analogous preparatory markers for particular movements would provide compelling evidence that infants’ action becomes intentional.

The very first movement emerges spontaneously during the fetal period, reflecting the endogenous activity of the developing nervous system (*7, 8*). Although the early spontaneous movements are not accompanied by prediction, infants gradually develop the ability to predict the consequences of their own actions. Piaget noticed the importance of the physical interaction with the environment for the transition from spontaneous movement to intentional action (*9*). His observations later led to the establishment of an experimental setup known as the mobile paradigm (*10*), in which infants’ arm or leg was tethered to a suspended mobile toy. This paradigm has been widely used to reveal the learning and memory capabilities of 2-to 3-month-old infants (*10–14*). It has been suggested that 3-month-old infants can perceive the causal relationship between their own movements and contingent mobile motion (*14–18*). EEG studies reported that neural components corresponding to mismatch negativity (*19*) or event-related potentials (*20*) were observed in 3-month-old infants, depending on whether their own movements caused the mobile to move. Although these studies provided the evidence of comprehension for action outcomes, they focused on the post-movement or post-stimulus component of neural activities. It is still unclear whether 3-month-old infants can initiate actions with prediction at self-determined timing. To ascertain whether intention emerges in 3-month-old infants interacting with their environments, it is necessary to measure some preparatory and predictive changes that occur before movement onset.

In the present study, we utilized changes in pupil diameter as an index of preverbal infants’ internal states, as the saying goes “the eyes are the window to the soul.” Pupil diameter is regulated not only by the autonomic nervous system but also by cortical processes, reflecting internal decision making (*21*) or emerging intention, similar to readiness potentials (*22*). Pupillometry is non-contact and resistant to movement artifacts, enabling more rigorous measurement than EEG in awake infants. Infants’ pupillary responses to visual images are comparable to adults’ one and have revealed wide range of infant’s cognitive capability, such as prediction and logical reasoning (*23–25*). However, no study has reported the pupil dynamics in relation to the timing of self-initiated movements in infants. Thus, in this study, we combined mobile paradigm and pupillometry to assess whether 3-month-old infants can initiate actions at self-determined timings with prediction for the action outcomes. To examine movement-related pupil changes, we first identified the timing of movement onsets using EMG signals. We then analyzed changes in pupil diameter before and after movement. Furthermore, we classified each movement according to EMG patterns and examined the relationship between movement types and changes in pupil diameter. In addition, we report gaze and heart rate changes during the mobile paradigm.

### Movement onset detection in freely moving infants

In our measurement, infants were lied on their back with two identical toys suspended above the infants (Fig. 1A). The two toys served as fixating points throughout the experiments, regardless of whether interaction occurred. The first 2-minute phase was a baseline phase when infants moved spontaneously with the toys suspended stationary. Subsequently, an infants’ arm and a toy ipsilateral to the arm were tied with an elastic string (Fig. 1B), initiating a 10-minutes interaction phase. The following phases were a 2-mimute extinction phase in which the physical interaction between the infant and the toy was removed and a 2-minute reinteraction phase in which the infant’s arm was again connected to the mobile (Fig. 1C). Twenty 3-month-old infants (12 girls and 8 boys, mean age 110.9±7.1 days) completed the whole procedure. To capture the onset of arm movements, we measured EMG signals from the deltoid, biceps, and triceps of the arm connected to the mobile toy (Fig. 1D).

**Fig. 1.**
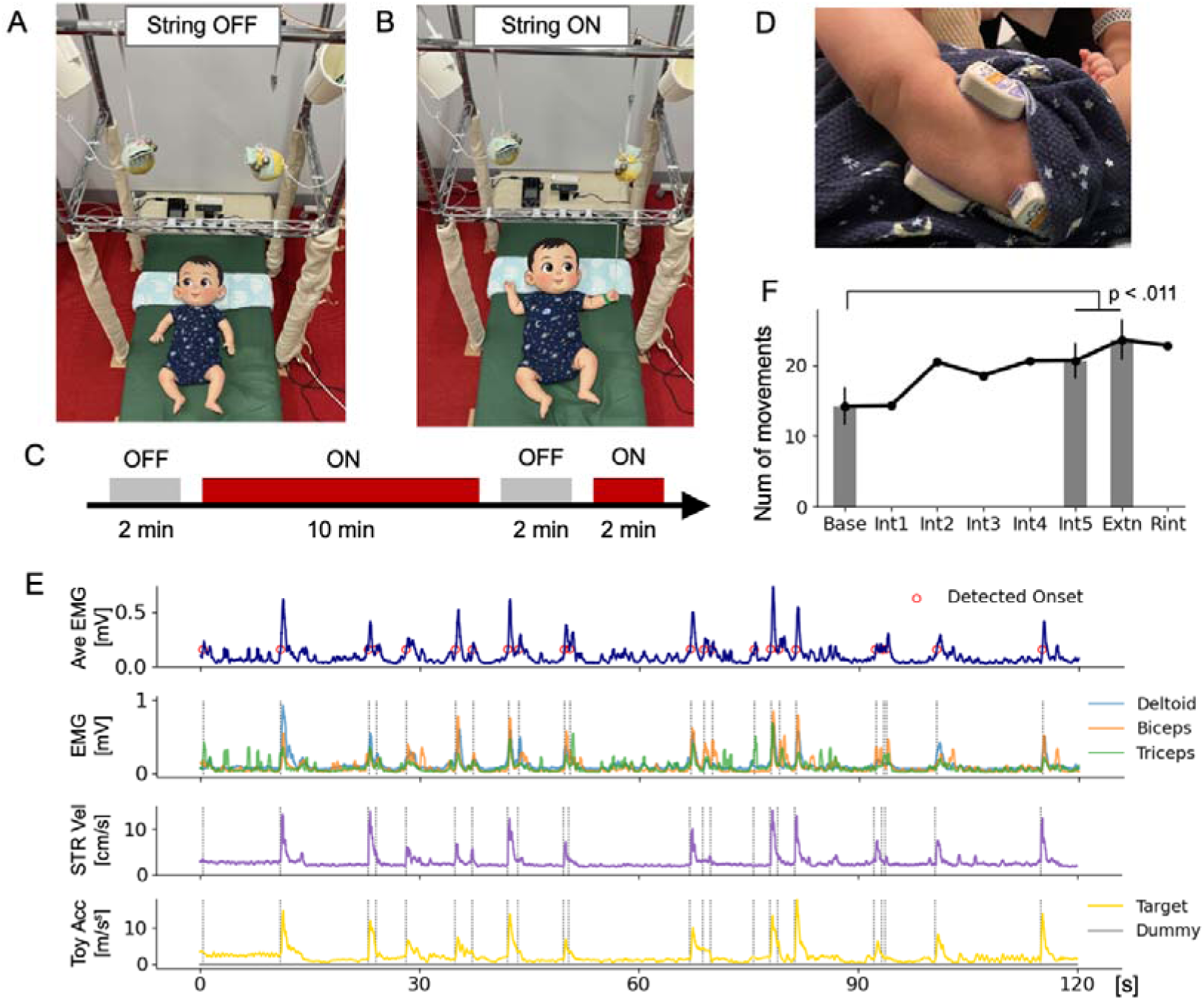
Onsets of arm movements were extracted using EMG. (**A**) An experimental phase without physical interaction between the infant and overhead mobiles. (**B**) The other experimental phase, in which the infant’s arm was connected to the overhead mobile via an elastic string. (**C**) The experimental procedure consists of four phases: a 2-min baseline phase without mobile connection, a 10-min interaction phase with mobile connection, a 2-min extinction phase without mobile connection, and a 2-min reinteraction phase with mobile connection. (**D**) Three EMG sensors were positioned on the deltoid, biceps, and triceps of the arm connected to the mobile. (**E**) An example of movement onset detection by using rectified EMG timeseries. We labelled the onset of movements when the averaged EMG exceed a threshold, which is differed from infant to infant, for more than 300 ms. The string was stretched (purple) and the connected mobile was accelerated (yellow) following the movement onset. (**F**) The time variation in the number of detected movements (Base: baseline phase, Int: interaction phase, Extn: extinction phase, and Rint: reinteraction phase). Error bars represent SEM across infants. The number of movements increased significantly after the interaction with the mobile (two-sided paired *t* test with Bonferroni correction after one-way repeated measured ANOVA, *n* = 20).

Arm movements were defined to occur when the averaged EMG signals exceeded a threshold (Fig. 1E), which was determined for each infant comparing EMG signals and actual arm movements through video. Immediately after the detected onsets, we observed peaks in the stretch velocities of the string (purple) and the acceleration of the connected toy (yellow), which validates our detection method (Fig. 1E). By using EMG signals, movement onsets were precisely identified from freely moving infants. The number of detected movements increased throughout the measurement (Fig. 1F). One-way repeated measures analysis of variance (ANOVA) confirmed that the number of detected movements differed across phases (*F*(7, 133) = 3.16, *P* = 4.1×10^-3^) and post-hoc two-sided paired *t* tests with Bonferroni correction confirmed that the number of movements significantly increased after the interaction with the mobile toy (Fig. 1F; Base vs Int5: *P* = 1.07×10^-2^, Base vs Extn: *P* = 2.95×10^-3^). Thus, our method replicated the increase in the number of movements, which has been used as a general indicator of learning in the mobile paradigm (*10–15*). Similar time evolutions were also observed in 2-min averaged heart rates during the measurements (fig. S1). Individual plots for the number of movements and heart rates further demonstrate that various patterns in infants’ behavioral changes were correlated with their heart rate changes (fig. S2-S5). The detected timings of movement onsets were subsequently used for analysing movement related pupil dilations.

### Gaze and pupil size analysis during the environmental interaction

In the history of developmental psychology, gaze durations during the mobile interaction were measured to assess infants’ attention (*9, 26, 27*). In recent years, eye-tracking systems have become a common method for measuring infants’ gaze toward the interactive toys or videos (*28–31*) and for pupillometry (*25, 32, 33*). In the present study, we utilized Tobii pro nano (Tobii Technology, Stockholm, Sweden) near-infrared gaze tracking system to enable the rigorous measurement of gaze points and pupil diameters at 60 Hz during the environmental interaction. Furthermore, we used spherical toys, making it easier to identify the gazing at the toy, and added another toy which infants could not interact with, allowing the comparison of gaze duration according to the presence or absence of physical interaction (Fig. 2A). Hereafter, the toy which infants interacted with is called the target toy and the other is called the dummy toy. During the measurement, we turned off the fluorescent light above the experimental field and covered the experimental setup with plain fabric to reduce the light-induced pupil constriction. The toys were illuminated by light outside the infants’ field of view.

Because infants mostly gazed at the target toy during the interaction phase, we defined that infants are gazing at the toy when their gaze point was within the area of concentrated gaze (Fig. 2B). The gaze duration at the target and dummy toy was then calculated for each 2-minute window (Fig. 2C). Two-way repeated measured ANOVA (2 toys × 8 phases) and post-hoc two-sided paired *t* tests (*n* = 20) with Bonferroni correction confirmed that infants gazed at the target toy significantly longer than the dummy toy during the interaction or reinteraction phase. The gaze durations for the target toy in these phases were significantly longer than those in the baseline and extinction phase, while the gaze duration became relatively shorter in the reinteraction phase (see fig. S6 for statistical details). These statistical significances reflect the shift in the infants’ attention toward the moving target toy and habituation for the toy movements. We also calculated the average pupil diameter within each 2-minute window (Fig. 2D). One-way repeated measured ANOVA revealed that the averaged pupil diameter changes across the phases (*F*(7, 133) =2.7, *P* = 1.2×10^-2^). Subsequent two-sided paired *t* tests with Bonferroni correction confirmed that the pupil diameter significantly shrank during the interaction phase (Int1 vs Int3: *P* = 1.73×10^-3^, vs Int4: *P* = 2.0×10^-4^). The pupil dynamics in the interaction phase may also reflect the shift in infants’ attention and the habituation for the environmental change. Particularly in the Int1 phase, pupils relatively dilated while their heart rate decreased (fig.S1), suggesting the occurrence of orienting responses (*34, 35*). Individual plots for gaze duration and pupil diameter demonstrate the differences and similarities across participants (fig. S7, S8).

**Fig. 2.**
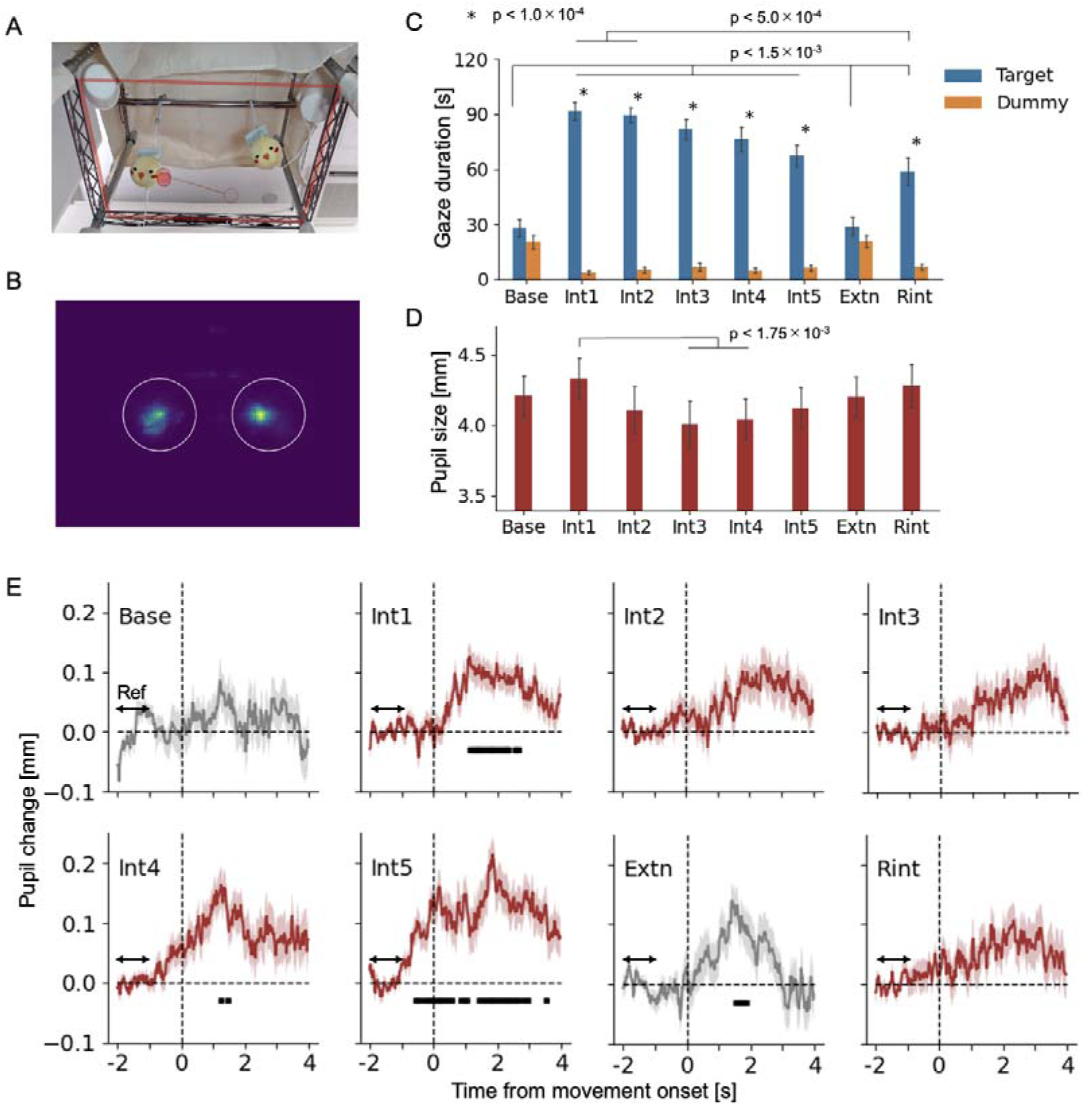
Infants gazed at the toy when they moved it, and pupil dilations came to precede the movement onset. (**A**) An example of gazing at the target toy. The red lines highlight the virtual plane defined for the gaze analysis. (**B**) The distribution of gaze points of all 20 infants during the interaction phase. We defined that infants were gazing at the target or dummy toy when their gaze points were in the white circles. (**C**) The averaged gaze duration for the target and dummy toy. Error bars are SEM across infants. Infants gazed at the target toy significantly longer than the dummy toy when they moved it (simple main effect in two-way ANOVA). The gaze duration for the target toy in Int1-5 and Rint phase were significantly longer than that in the Base and Extn phase. In the Rint phase, the gaze duration was significantly shorter than that in the Int1 and Int 2 phase (two-sided paired *t* test with Bonferroni correction, *n* = 20). (**D**) The averaged pupil diameter in each phase. Error bars are SEM across infants. The pupil diameter in the Int1 phase was significantly larger than that in the Int3 and Int4 phases (two-sided paired *t* test with Bonferroni correction, *n* = 20). Error bars represent SEM across infants. (**E**) Pupil dilations associated with the onset of arm movements. Shadow areas are SEM across trials. ‘Ref’ indicates the interval used as a baseline pupil diameter whose average was adjusted to zero in each trial. Black bars indicate intervals of significant pupil dilation (*P* < 0.05) by linear mixed model and one-sample one-sided *t* test (versus 0), applied every 0.1-second interval with sequential Bonferroni. Pupil dilation associated with movement emerged in Int1 phase, and gradually shifted to precede movement onset in Int4 and Int5 phase. Significant pupil dilation was also observed in Extn phase.

### Pupil dilation comes to precede the movement onset

We conducted further analysis of phasic pupil dilation, time-locked to movement onsets, focusing on trials in which infants maintained their gaze on the toy. We first extracted movements that occurred without any prior movements within a 6-second window to avoid potential contamination of pupil dilation caused by prior events. Furthermore, we selected the trials in which the measured rate of pupil diameter exceeded 50% and the rate of target gaze exceeded 65% (fig. S9) in the interval from 2-second before to 4-second after the movement onset (table. S1, S2). In the baseline and extinction phase, the rate of target gaze was adjusted to the summed rate of target and dummy gaze. For each trial, a baseline pupil diameter was computed by averaging the pupil diameters within the 1-second window starting 2 seconds before the movement onset. We subtracted the baseline pupil diameter from the raw pupil sizes in order to extract the changes in pupil dimeter associated with the movement onset and then averaged these changes within each phase (Fig. 2E). During the baseline phase, no pupil dilation was observed associated with the movement onset. However, once the infant’s arm was tethered to the target toy and the toy started to move (fig. S10), pupil dilation following arm movements came to appear. This trend continued into the middle of the interaction phase, although the statistical significance was not observed during the Int2 and Int3 phases. From the Int4 phase, the pupil dilation gradually began to precede the initiation of arm movements, and by Int5 phase, pupil dilation prior to arm movement was observed with statistical significance (Fig. 2E). Such changes would not occur unless infants possessed some form of awareness regarding the timing of their forthcoming arm movements. While the antecedent pupil dilation disappeared during the extinction phase, pupil dilation following arm movements persisted, even though the physical setup was identical to the baseline phase. In contrast, heart rates did not show such changes and instead exhibited a suppressed increase in the Int5 phase (fig. S11), inferring the dissociated control of pupil size related to the prediction for action outcomes. We further investigated the relationship between the baseline pupil diameter for each trial and the magnitude of subsequent pupil dilation (fig. S12, S13). Negative correlations were predominantly observed between the baseline pupil size and subsequent pupil dilation, indicating that phasic changes were modulated by tonic changes. However, the pre-movement dilation in Int5 phase and the post-movement dilation in the extinction phase did not exhibit significant negative correlations. These results suggest that pupil dilation associated with prediction involves mechanisms that are relatively independent of tonic pupil sizes.

### Alterations in the EMG patterns and their relation to antecedent pupil dilation

Intended movements may differ in their efficiency and magnitude from those occurred without preparations in terms of muscular activities. Thus, it is possible that EMG coordination patterns are related to the occurrence of antecedent pupil dilations. To reveal the hidden strategies of infants’ arm movements during the interaction with the mobile, we applied time-series clustering to the EMG signals of three muscles within 500 ms after each movement onset. As a result, the EMG signals were divided into five clusters (Fig. 3A and B). The pattern in the largest cluster showed small amplitude activity across the three muscles. The following three clusters consist of high amplitude activity in each single muscle: deltoid, triceps, and biceps. The smallest cluster contains movements with high amplitude activity in all three muscles, particularly in deltoid and triceps.

**Fig. 3.**
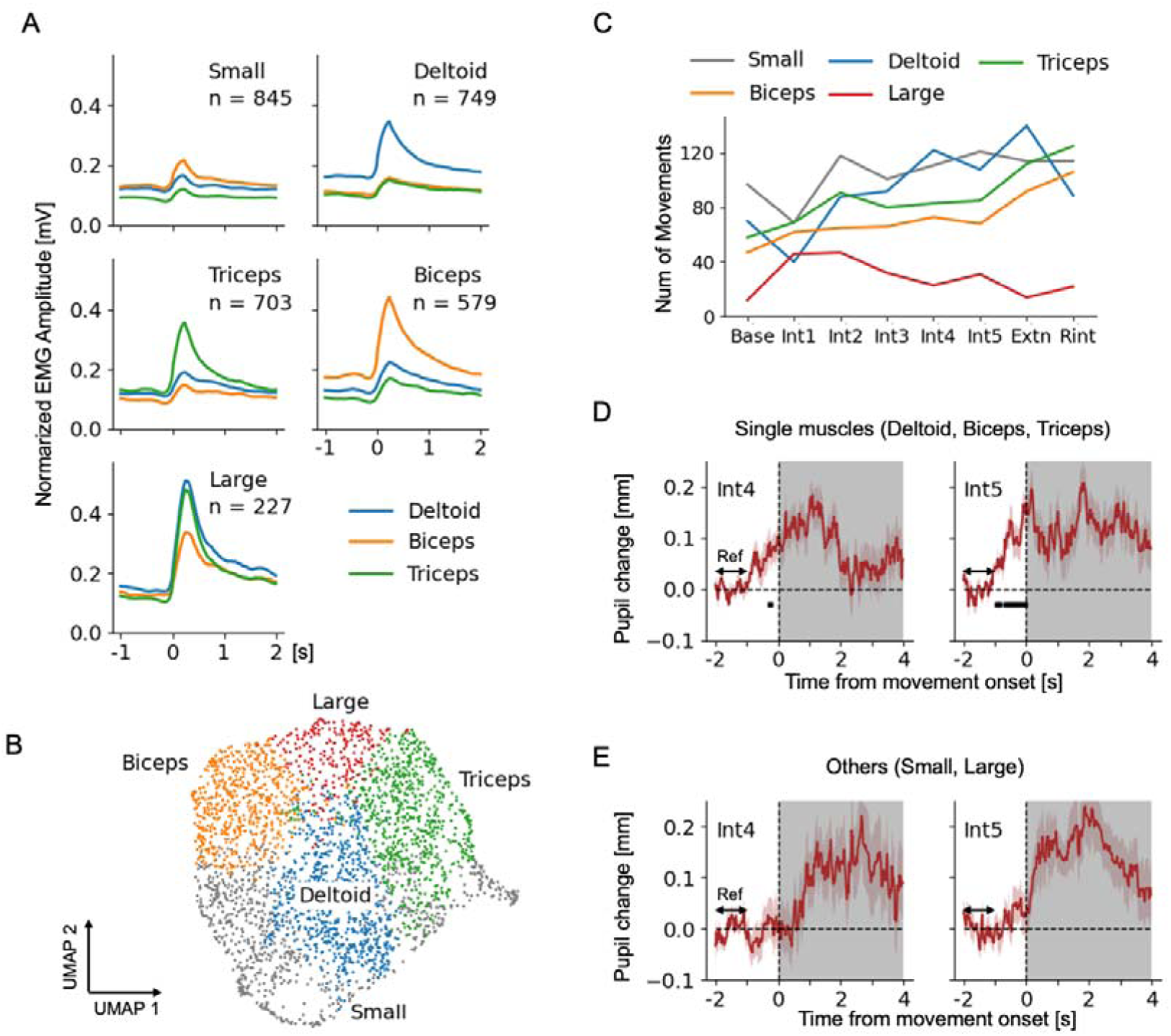
EMG patterns of arm movements associated with predictive pupil dilation. (**A**) EMG activity of three muscles around movement onset were divided into five subgroups using time-series clustering. Colors indicate the muscles. (**B**) UMAP embedding of three-muscle EMG patterns between 1-seconds before and 2-seconds after the movement onsets. The colors and labels represent the clusters, not muscles. (**C**) The time evolutions in the number of movements in each cluster, summed across infants. The colors and labels represent the clusters, not muscles. (**D** and **E**) Pupil dilations associated with the onset of arm movements: (D) movements dominated by a single muscle, clustered as Deltoid, Biceps, or Triceps; and (E) movements clustered as Small or Large. Red shadow areas are SEM across trials. ‘Ref’ indicates the interval used as a baseline pupil diameter whose average was adjusted to zero in each trial. Black bars indicate intervals of significant pupil dilation (*P* < 0.05) by linear mixed model and one-sample one-sided *t* test (versus 0) with sequential Bonferroni, applied every 0.1 second during the interval preceding the movement onset. Antecedent pupil dilation was only observed in the single-muscle dominant group (D).

The total number of movements across infants in the baseline phase (Fig. 3C) indicates that spontaneous movements consist of many small movements, a few large movements, and a moderate number of movements involving single muscle activities. This composition changed drastically after the mobile connection with decrease in small movements and increase in large movements. Subsequently there was a rebound in the number of small movements and increase in the number of movements caused by single muscle activations. In contrast, the number of large movements decreased gradually during the interaction phase. There was substantial individual variation in which of these three muscles was primarily used. (fig. S14). To examine the relationship between EMG coordination patterns and pupil dilation regardless of this individual variation, the three single-muscle dominant clusters were grouped together as a single-muscle group (fig. S15). The association between antecedent pupil dilation and muscular activity patterns was analyzed by comparing this single-muscle group with the other movement group (small and large) (table. S3). As a result, the antecedent pupil dilation was observed only in the single-muscle group in both Int4 and Int5 phases (Fig. 3D, E), suggesting the relationship between the awareness for the forthcoming movement and the efficiency of executed actions.

### The antecedent pupil dilation reflects emerging intention to move

Combining the mobile paradigm with pupillometry, we found that pupil dilation comes to precede self-generated actions through environmental interactions. In previous studies using mobile paradigm, it has been argued that infants can learn the causal relationship between their own movements and the mobile motion (*14–18*). Recent studies also support the view that 3-month-old infants anticipate outcomes of their actions (*19, 20, 36*). However, these studies do not ascertain whether infants can initiate actions with prediction at self-determined timing, which is essential for intentional actions. In this study, we revealed the emergence of antecedent pupil dilation, which does not occur unless infants know the timing of their own actions in advance. Notably, pupil dilation associated with movement did not occur during the baseline phase, highlighting the importance of environmental interaction. Through the sensorimotor coordination, infants came to perceive not only the causal relationship between their movements and subsequent events, but also that they are the agents of movements, enabling action generation with prediction.

When investigating the link between antecedent pupil dilations and EMG patterns, we found that only arm movements with single-muscle dominant activity were accompanied by antecedent pupil dilations. Single muscle activities are considered to be more efficient in producing arm movements and may reflect the preparation for action generation. In contrast, antecedent pupil dilation was not observed when muscle activity was minimal or all three muscles contracted simultaneously. These movements may occur without prediction and could be an extension of spontaneous movements. Thus, it is possible that intentional and unintentional movements coexist — sometimes infants move their bodies intentionally, and other times their body moves spontaneously.

The antecedent pupil dilation has been reported to occur in human adults prior to their voluntary action or decision making (*21, 22*), and discussed in relation to the readiness potential (*22*). Pupil diameters are mainly regulated by the locus coeruleus and the superior colliculus under constant lighting conditions, reflecting arousal levels and shifts in attention (*34, 35, 37*). However, pupil diameters are also influenced by cortical activities with particular involvement of the medial prefrontal cortex (mPFC) (*22, 38, 39*). During the first three months, the inter-cortical and thalamocortical circuits mature via the subplate, enabling adult-like neural activities (*40–42*). It has been shown that the mPFC is functionally active in 3-month-old infants while they perceive audiovisual stimuli from mobile toys (*43, 44*). In adult, the mPFC is also known to play a crucial role in initiating voluntary actions (*45*) and predicting action outcomes (*46*). Thus, it is possible that the mPFC, already functional at three months, plays a role in initiating actions with outcome predictions, serving as a foundation for the formation of intentional action.

In this study, we captured a physiological marker that reflects emerging intention through the environmental interaction. However, our results do not necessarily suggest that intention itself first emerged at this moment. It is also possible that intention already exists latently and becomes apparent through interaction with the environment. To clarify the emergence of intention itself, longitudinal studies are needed starting from younger ages. For example, newborn infants exhibit evidence of reinforcement in non-nutritive sucking with the mother’s voice (*47*), arm movements with facial images of mothers (*48*), and arm movements with the live video of their own arm movements (*49*). It is also reported that there is a developmental change from newborns to 2-month-old infants in sucking behaviors with contingent sounds (*50*). By combining such experimental paradigms with pupillometry, it may be possible to capture the emergence of intention from the neonatal period. Our findings pave the way for assessing intention, by measuring pupil dynamics while infants actively interact with their environment.

## Materials and Methods

### Participants

Twenty 3-month-old infants completed the experimental procedures (12 girls and 8 boys, age 99-127 days, mean age 110.9±7.1 days). Additional eight infants participated in the present study but were excluded from the sample because of crying (*n* = 5) and rolling over (*n* = 3). All participants were full-term infants recruited via the local Basic Resident Register. Ethical approval for this study was obtained from the ethical committee of Life Science Research Ethics and Safety, the University of Tokyo, and written informed consent was obtained from the parent(s) of all the infants prior to the initiation of the experiments.

### Apparatus

In the current study, two identical non-commercial hand-made toys were used as mobiles, one for a target toy and the other for a dummy toy. Both toys were suspended above infants during the measurement but only the target toy was tethered to an infant’s arm during interaction phases. The dummy toy served as another fixating point during the experiment. The selection of the target toy, left or right, was counterbalanced between participants. The toy was made of a soft cotton ball covered by a sock which contains a face of yellow chick. Two small bells were attached to each toy leading to make attractive sounds when the toy was moved. During the measurement, each infant was positioned on their back on a baby mattress (width: 55cm, depth: 110cm) and the toys were suspended above the infant from a pole passed at the top of a stainless frame (width: 60cm, depth: 45cm, height: 120cm). The stainless frame consisted of four 120cm-height poles and a U-shaped frame with right-angle corners (width: 60cm, depth: 45cm). The U-shaped frame was located 70cm above the baby mattress and utilized as a virtual plane for eye tracking system. The vertical positions of toys were kept on the virtual plane, using hard elastic strings for suspending toys and a soft elastic string for connecting the target toy and an infant’s arm.

In this setting, we measured eye movements, pupil diameters, electrocardiogram (ECG), and electromyogram (EMG) of the connected arm (deltoid, biceps, and triceps) from infants; and measured accelerations of each toy and stretches of the string suspending the target toy. Additionally, a video camera was used to record other information such as facial expressions, gross movements, and the status of the mobile. Infant’s eye movements and pupil diameters were estimated using Tobii pro nano (Tobii Technology, Stockholm, Sweden) near-infrared gaze tracking system, which records near-infrared reflections of eyes at 60 Hz while infants were watching objects within the predefined virtual plane. ECG, EMG of the connected arm, and accelerations of each toy were measured at 1000 Hz via multi-channel telemeter system WEB-1000 (Nihon Kohden, Tokyo, Japan). Stretches of the string were measured by C-Stretch measure (Bando Chemical Industries, Hyogo, Japan) and recorded at 1000 Hz via multi-channel telemeter system.

### Procedures

All measurements were conducted in the laboratory at the Graduate School of Education of the University of Tokyo. Prior to the experiment, infants were laid on a mattress out of the test field and were attached EMG and ECG sensors. As the infant were likely to alert and playful, an experimenter carried the infant to the test field and measurements were carried out. At first, a calibration for the eye tracking system was carried out with a different toy inducing infant’s fixation, while the target and dummy toys were hidden in baskets. Immediately after the calibration, two experimenters moved the toys silently above the infant and the experiment started. Our experiment consists of four phases: 2-min baseline, 10-min interaction, 2-min extinction, and 2-min re-interaction. In the first 2-min baseline phase, infants moved spontaneously and did not cause the mobile to move, observing the motionless mobiles suspended above. After the baseline phase, a string was attached between the target toy and an infant’s wrist (the connected wrist was counterbalanced because the same side of the target toy was connected). This arrangement initiated an interaction phase lasting 10 minutes. During the interaction phase, any movements of the connected arm produced a corresponding degree of motion and sound of the target toy. After the interaction phase, the string was detached from the infant’s wrist and 2-min extinction phase was initiated, in which the infant could not make any movements of the target toy. At last, the string was again connected to the infant’s wrist as 2-min re-interaction phase.

### Data Processing

EMG data were applied moving root mean square (RMS) with 250ms time window to calculate the absolute values and to low pass filter the raw EMG time series. The data of stretches of the string were differentiated to calculate their change velocity and then applied moving RMS with 250ms time window to calculate the absolute values and low pass filter the time series. The data of accelerations of toys in a 3D space (x, y, z) were first detrended to remove the effect of the gravitational acceleration. Subsequently, the RMS of the data in three axes (x, y, z) were calculated, and then applied moving average with 250ms time window to low pass filter the time series. ECG data ware converted into R-wave intervals and time series of heart rate by using the Kubios HRV Analysis Software 2.0 (The Biomedical Signal and Medical Imaging Analysis Group, Department of Applied Physics, University of Kuopio, Finland) (*51*). The ECG data from one participant could not be reliably converted into heart rate and was excluded from subsequent analysis.

### Movement onset detection

We detected onsets of the arm movements by using rectified EMG timeseries. Each rectified EMG data for three muscles were normalized to the maximum value of 1 and then averaged. We applied thresholds, differed from infant to infant, to the averaged EMG timeseries to detect the onset of movements. The thresholds were determined as 8 to 18 % of the range between maximum and minimum of the peak values in the averaged EMG timeseries. If the averaged EMG timeseries exceeds this threshold more than 300 ms, we defined the onset of the section as a movement onset.

### Time series clustering

We conducted the time-series clustering to elucidate the various patterns of muscle coordination for the arm movements to move the mobile. This method divides the time-series data into a certain number of groups by calculating its similarities, which utilized to elucidate the hidden strategies in infants’ behaviours (*52*). We used K-means time-series clustering with Euclidian distance by adopting the following equation:

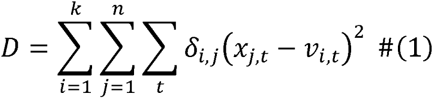

Assuming that there are *n* time-series data {*x_j,t_*│*j*=1,2,⋯,*n*} which are to be divided into *k* clusters. The number of clusters, *k*, is a parameter decided to capture the structure of data best. In our study, *k* was decided as *k* = 5. *D* represents the sum of distance between each time-series data from their respective cluster centers {*v_i,t_*│*i*=1,⋯,*k*}. *δ_i,j_* = 1 if *x_j,t_* is assigned to cluster *i*, and otherwise *δ_i,j_* = 0. The main idea of K-means clustering is the minimization of the function *D* by altering the assignment of each time-series data to the appropriate cluster (*53*). At first, each data was randomly allocated to one of the clusters and then, iteratively change its allocation to the cluster whose center is nearest to the time-series data until the function *D* cannot be reduced anymore. In our study, *x_j,t_* represents the time-series of rectified EMG amplitude of three muscles within 500 ms after each movement onset.

The function was implemented by using tslearn (*54*), a Python machine learning package for time series data. tslearn depends on basic Python packages, NumPy and SciPy, for array manipulation and standard linear algebra routines, and follows scikit-learn’s Application Programming Interface, which integrates a wide range of machine learning algorithms for Python.

### Gaze analysis

For gaze analysis, the U-shaped frame (width: 60cm, depth: 45cm) located 70cm above the baby mattress was utilized as a virtual plane. This virtual plane was resized to 1080×1920 pixels and gaze points were recorded on this plane. Because infants mostly gazed at the target toy during the interaction phase, we defined that infants are gazing at a toy when the infants’ gaze point was in the area where the gaze points were concentrated. We defined two circles with a 400 pixels diameter whose center was located at (670, 650) or (670, 1250) on the virtual plane as the gaze-concentrated areas.

### Preprocessing for time series of pupil diameters

To remove outliers, we first excluded instances where the difference between two consecutive data points exceeded 0.2. Subsequently, we applied a five-point moving average to reduce noise.

### Preprocessing for analysing pupil change with movements

We first identified movement onsets that occurred without any prior movement onsets within a 6-second window to avoid potential contamination of pupil dilation caused by prior events. For each of these onsets, we extracted the timeseries of right and left pupil diameters from 2 seconds before to 4 seconds after the movement onset, and then averaged the right and left pupil diameters in each time point. Data exceeding ±3 standard deviations from the mean were additionally removed at this stage. We selected the trials in which the measured rate of pupil diameter exceeded 50% (Fig. 3E) and the rate of target gaze in the measured gaze points exceeded 65% (Fig. 3F). In the baseline and extinction phase, the rate of target gaze was adjusted to the total rate of target and dummy gaze. The number of trials used for the pupil dilation analysis was 21 for baseline phase, 81 for interaction phase 1 (Int1: 0-2 minutes), 72 for Int2, 82 for Int3, 69 for Int4, 54 for Int5, 29 for extinction phase, and 56 for reinteraction phase. For each trial, a baseline pupil diameter was computed by averaging the samples within a 1-second window starting 2 seconds before the movement onset. Subsequently, we computed the difference between raw pupil diameters and the baseline pupil diameter in each time point to extract the changes in pupil dilation associated with the onset of arm movements.

### Statistical analysis

For the statistical analysis of the number of movements (Fig. 1F) and the averaged pupil diameters (Fig. 2D), we conducted one-way repeated measures analysis of variance (ANOVA) because phase was considered as a within-subject factor. For the analysis of the gaze duration at the target and dummy toy (Fig. 2C), we conducted two-way repeated measures ANOVA because both phase and toy were considered as within-subject factors. All ANOVA analysis was conducted using Python packages, NumPy and Pandas. We applied two-sided paired *t* tests with Bonferroni correction as post-hoc multiple comparisons for repeated measures ANOVA. *T* tests were implemented by using stats from a Python package SciPy. For the statistical analysis of the degree of pupil dilation related to movement onsets, we used a linear mixed-effects model (LMM). We estimated the degree of pupil dilation at 0.1-second intervals as a fixed effect while accounting for individual differences in baseline pupil diameter by including participant　as a random intercept. To examine whether the fixed effect was significantly larger than zero, we conducted one-sided *t* tests at every 0.1-second interval, with sequential Bonferroni procedures applied for the multiple testing corrections. We applied one-sided *t* tests because previous studies (21, 22) have shown that pupils dilate around movement onsets and our data contain only phasic pupil dilations throughout the entire phases. Python package MixedLM from statmodels were used for this analysis. *P* value threshold was 0.05 in our entire analysis.

## Supplementary Information

**Fig. S1.**
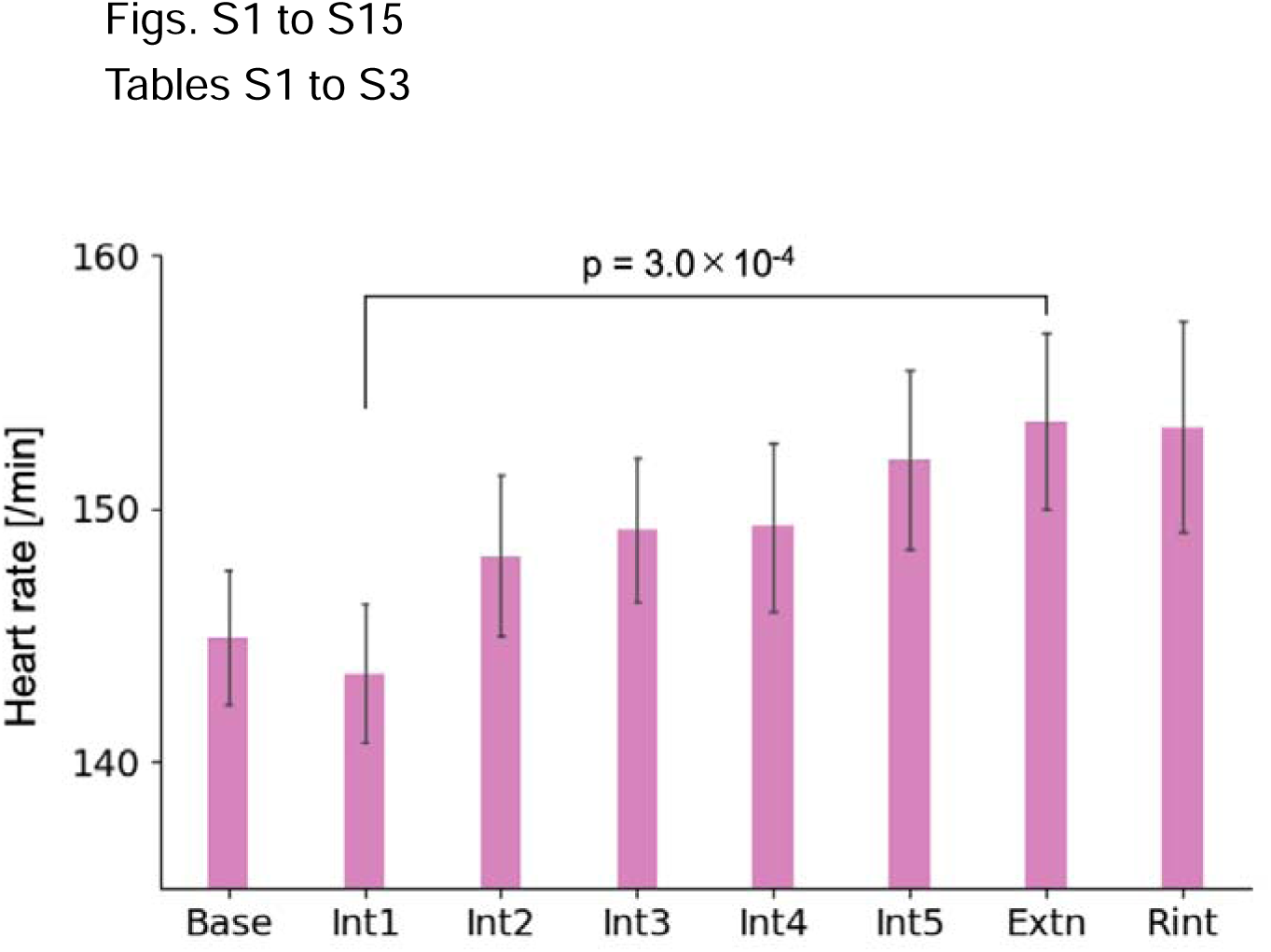
The averaged heart rate in each phase. Error bars are SEM across infants (*n* = 19). One-way repeated measured ANOVA revealed that the averaged heart rate changes across the phases (*F*(7, 126) = 4.4, *P* = 2.0×10^-4^). Subsequent two-sided paired *t* tests with Bonferroni correction confirmed a significant increase in heart rate from Int1 to Extn phase (*P* = 3.0×10^-4^). The time course of averaged heart rate was comparable to that of movement frequency (Fig. 1F). Additionally, the decrease in heart rate during the Int1 phase, coinciding with pupil dilation (Fig. 2D), suggests the occurrence of orienting responses.

**Fig. S2.**
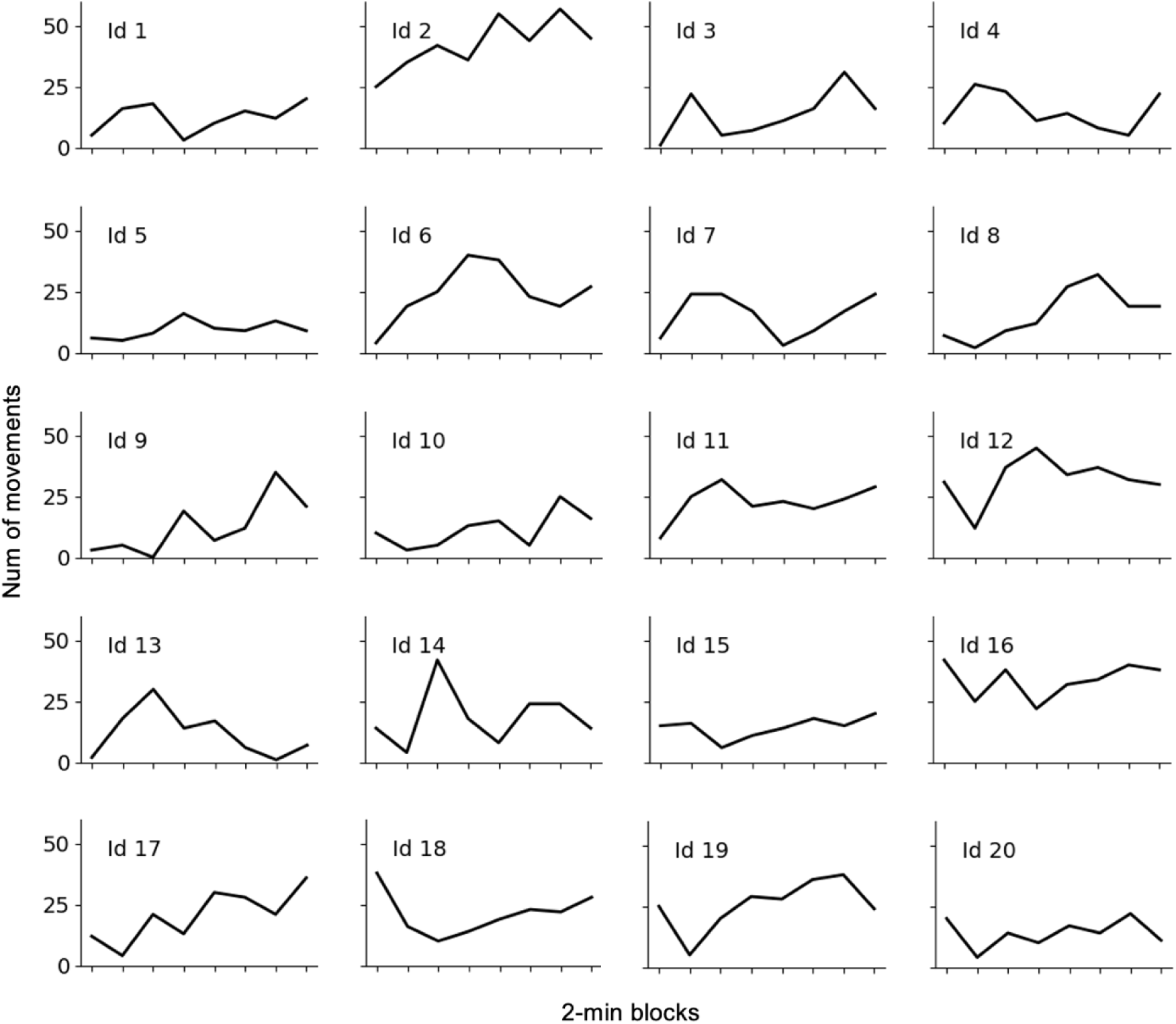
Individual plots for the number of movements in 2-min intervals. The marks on the X-axis correspond, from left to right, to Base, Int1-Int5, Extn, and Rint. There is a wide variety of patterns in the total number of movements, the degree of increase during the interaction, and the baseline level. A number of infants (Id 12, 14, 16, 17, 18, 19, 20) exhibited a clear decrease in movements during the Int 1 phase, while other infants (Id 1, 3, 4, 6, 7, 11, 13) showed an increase in movements in the same phase. This dissociation may be related to the frequency of spontaneous movements during the baseline phase. There were individual differences also in the changes of movement frequency during the extinction phase: Some infants increased their movements (Id 2, 3, 9, 10, 20), while others decreased them (Id 1, 4, 8, 17).

**Fig. S3.**
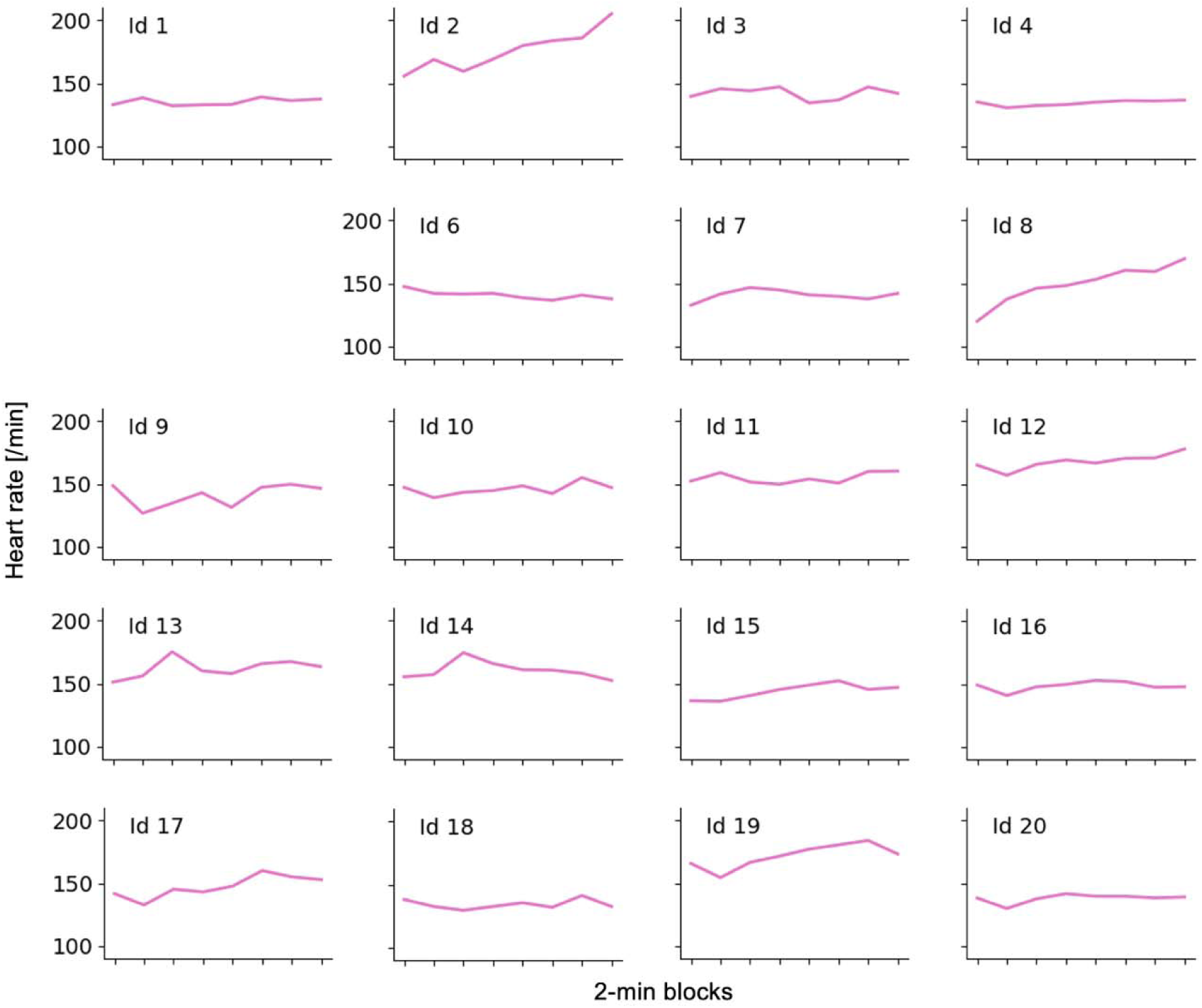
Individual plots for the 2-min averaged heart rate. The marks on the X-axis correspond, from left to right, to Base, Int1-Int5, Extn, and Rint. Inter-individual differences appear to be greater than intra-individual variabilities. Due to the system error, we omitted the data from an infant (ID 5). Two infants (Id 2 and Id 8) exhibited a certain increase in heart rate, which was due to mild fussiness observed in the latter part of the measurement. However, since they were still able to complete the task, their data were included in our analysis.

**Fig. S4.**
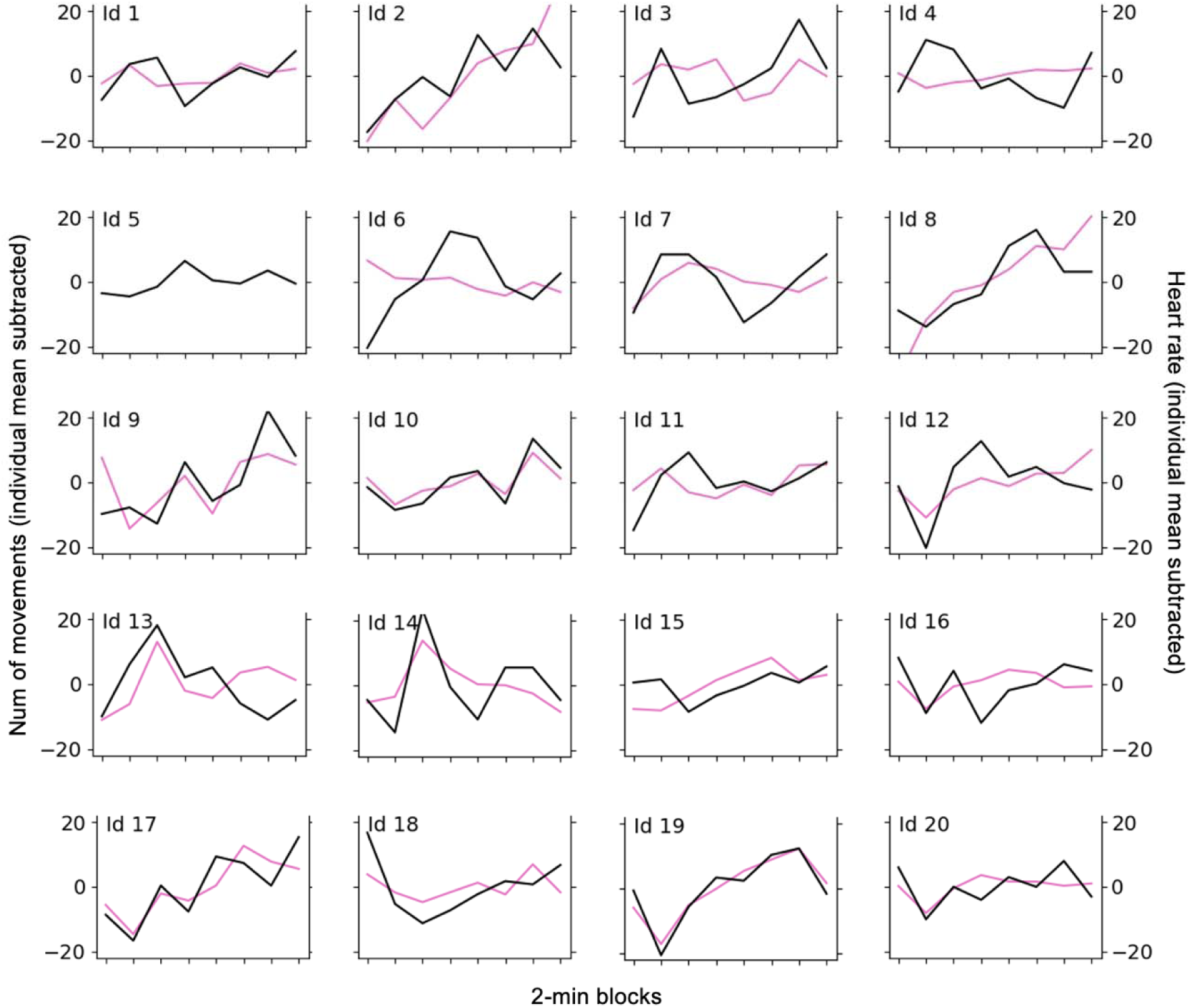
Time courses of the changes in the number of movements and heart rates in 2-min intervals. The marks on the X-axis correspond, from left to right, to Base, Int1-Int5, Extn, and Rint. The Y-axis represents the change from each individual’s average. There is a general tendency that changes in movement frequency is coupled to heart rate variability. Some infants (Id 10, 17, 19) exhibited almost the same changes in the number of movements and heart rate.

**Fig. S5.**
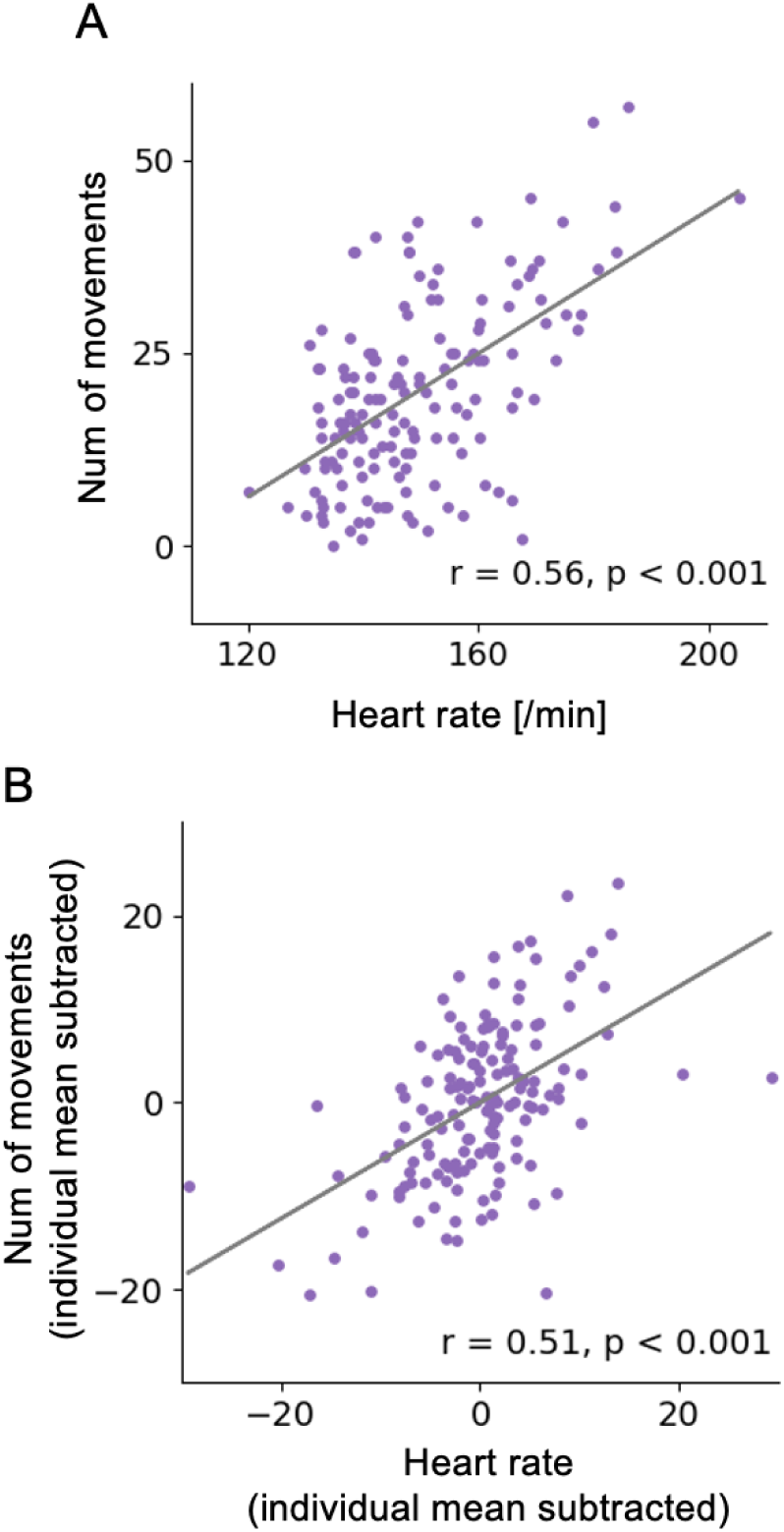
Correlation analysis between the number of movements and heart rate. One data point represents the value in each 2-min interval, thus, there are 8 data points per infant. (A) There is a moderate positive correlation (r = 0.56) between the number of movements and the heart rate in their raw values. (B) A moderate positive correlation (r = 0.51) was also observed between the number of movements and the heart rate in terms of their deviation from each individual’s average.

**Fig. S6.**
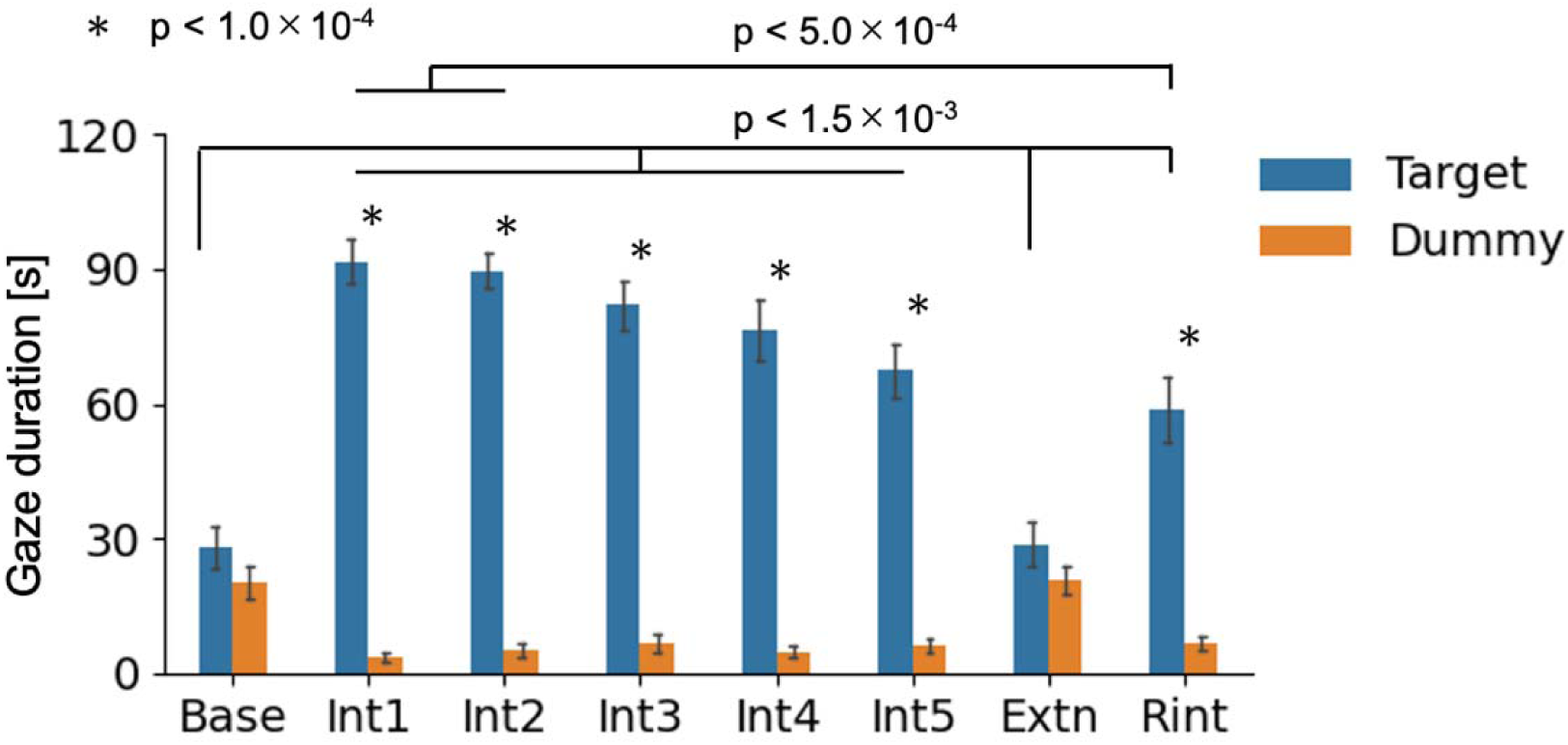
Statistical details for gaze duration. This figure is the same as Fig. 2C. The following descriptions are the statistical details. Two-way repeated measured ANOVA (2 toys × 8 phases) confirmed the significant main effects for the toys (*F*(1, 19) = 225.6, *P* = 5.4×10^-12^) and for the phases (*F*(7, 133) = 17.5, *P* = 3.3×10^-16^), and the significant interaction between the toys and phases (*F*(7, 133) = 32.4, *P* = 1.1×10^-16^). With regard to the interaction, there were significant simple main effects of the toys in the phases when infants interacted with the target toy (Int1: *F*(1, 19) = 159.2, *P* = 1.1×10^-10^; Int2: *F*(1, 19) = 192.4, *P* = 2.1×10^-11^; Int3: *F*(1, 19) = 78.1, *P* = 3.7×10^-8^; Int4: *F*(1, 19) = 53.5, *P* = 6.2×10^-7^; Int5: *F*(1, 19) = 46.3, *P* = 1.7×10^-6^; and Rint: *F*(1, 19) =24.0, *P* = 9.9×10^-5^). There were also significant simple main effects of the phases in the gaze duration at the target toy (*F*(7, 133) = 17.3, *P* = 3.3×10^-16^) and the dummy toy (*F*(7, 133) =8.7, *P* = 9.4×10^-9^). With respect to the target toy, post-hoc two-sided paired *t* tests with Bonferroni correction revealed that the gaze duration in the interaction and reinteraction phases was significantly longer than that in the baseline or extinction phases (Base vs Int1: *P* = 9.0×10^-12^, vs Int2: *P* = 2.2×10^-12^, vs Int3: *P* = 5.3×10^-9^, vs Int4: *P* = 6.3×10^-7^, vs Int5: *P* = 6.9×10^-6^, vs Rint: *P* = 8.6×10^-4^; Extn vs Int1: *P* = 3.5×10^-11^, vs Int2: *P* = 1.1×10^-11^, vs Int3: *P* = 1.5×10^-8^, vs Int4: *P* = 1.4×10^-6^, vs Int5: *P* = 1.5×10^-5^, vs Rint: *P* = 1.4×10^-3^), and the gaze duration in the reinteraction phase was significantly shorter than that in the Int1 (*P* = 4.0×10^-4^) and Int 2 phase (*P* = 4.9×10^-4^).

**Fig. S7.**
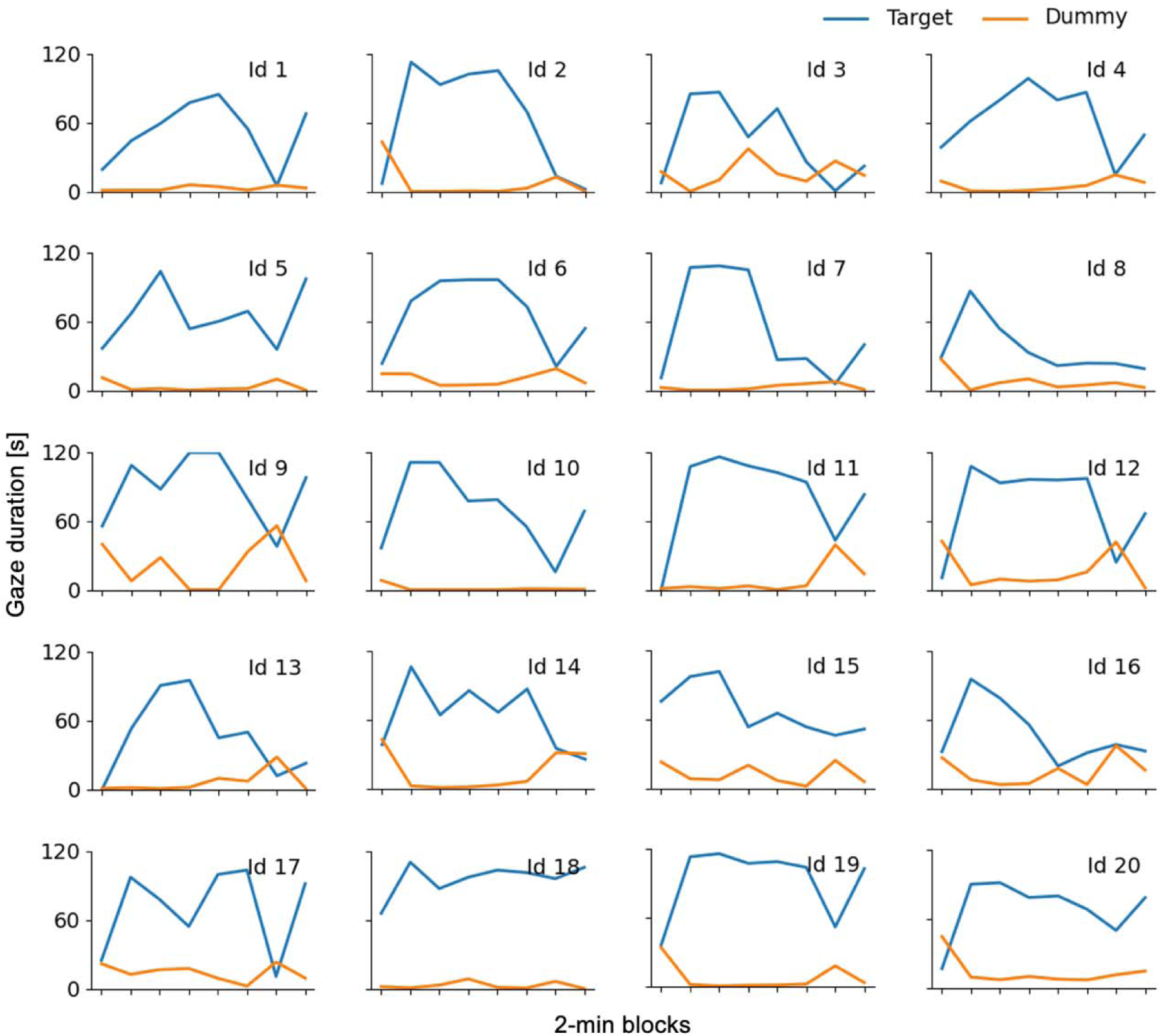
Individual plots for the gaze duration at the target toy and dummy toy. The marks on the X-axis correspond, from left to right, to Base, Int1-Int5, Extn, and Rint. There is a general tendency for infants to gaze at the target toy in the interaction phase, even at the individual level. While the decrease in gaze duration in two infants (Id 2 and Id 8) were due to mild fussiness, the decrease in other infants (Id3, 7, 10, 13, 15, 16) was caused by looking away, which may indicate habituation.

**Fig. S8.**
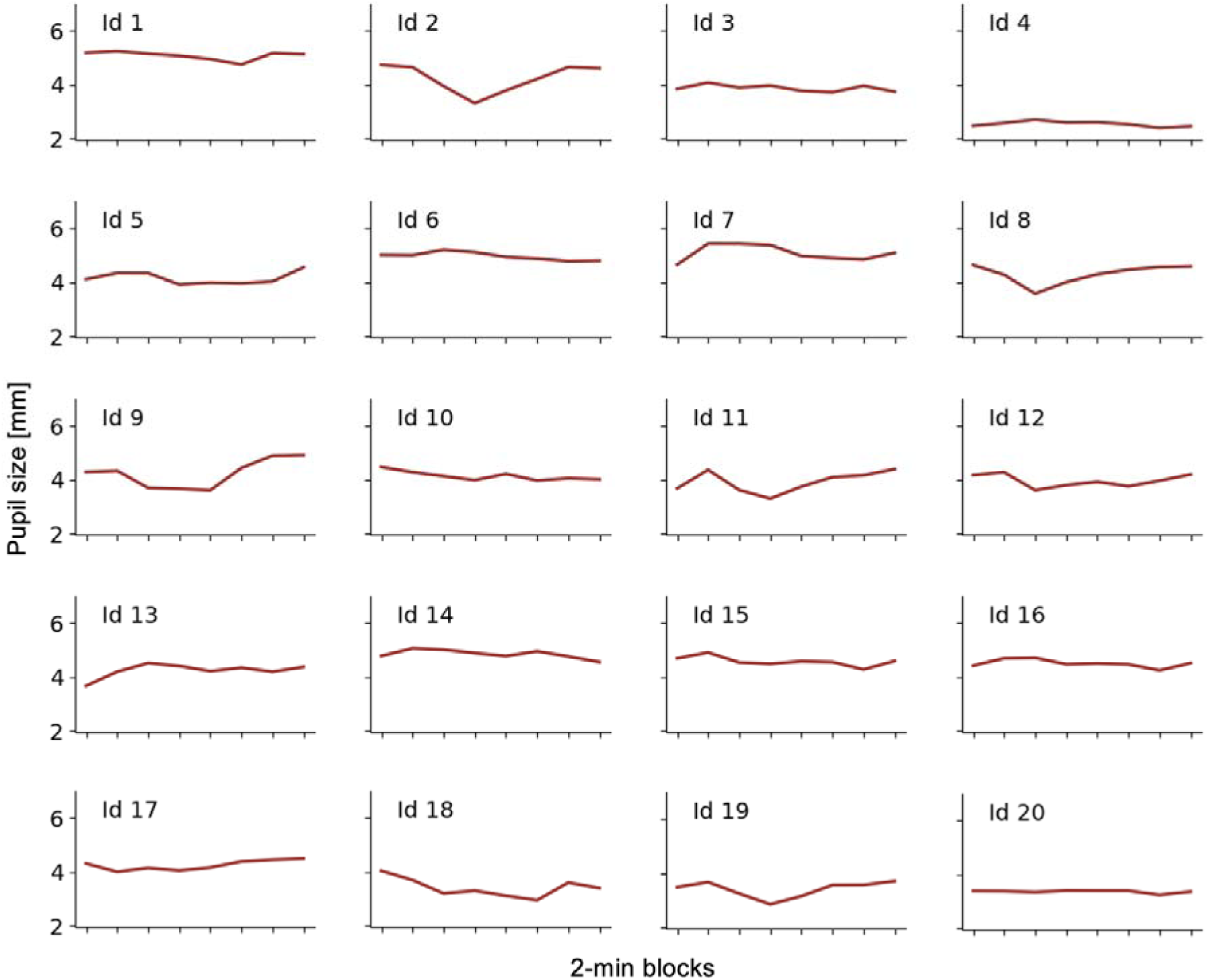
Individual plots for the 2-min averaged pupil diameter. The marks on the X-axis correspond, from left to right, to Base, Int1-Int5, Extn, and Rint. Inter-individual differences appear to be greater than intra-individual variabilities. Infants who exhibited mild fussiness (Id 2 and Id 8) showed a decrease in pupil diameter followed by a rebound.

**Fig. S9.**
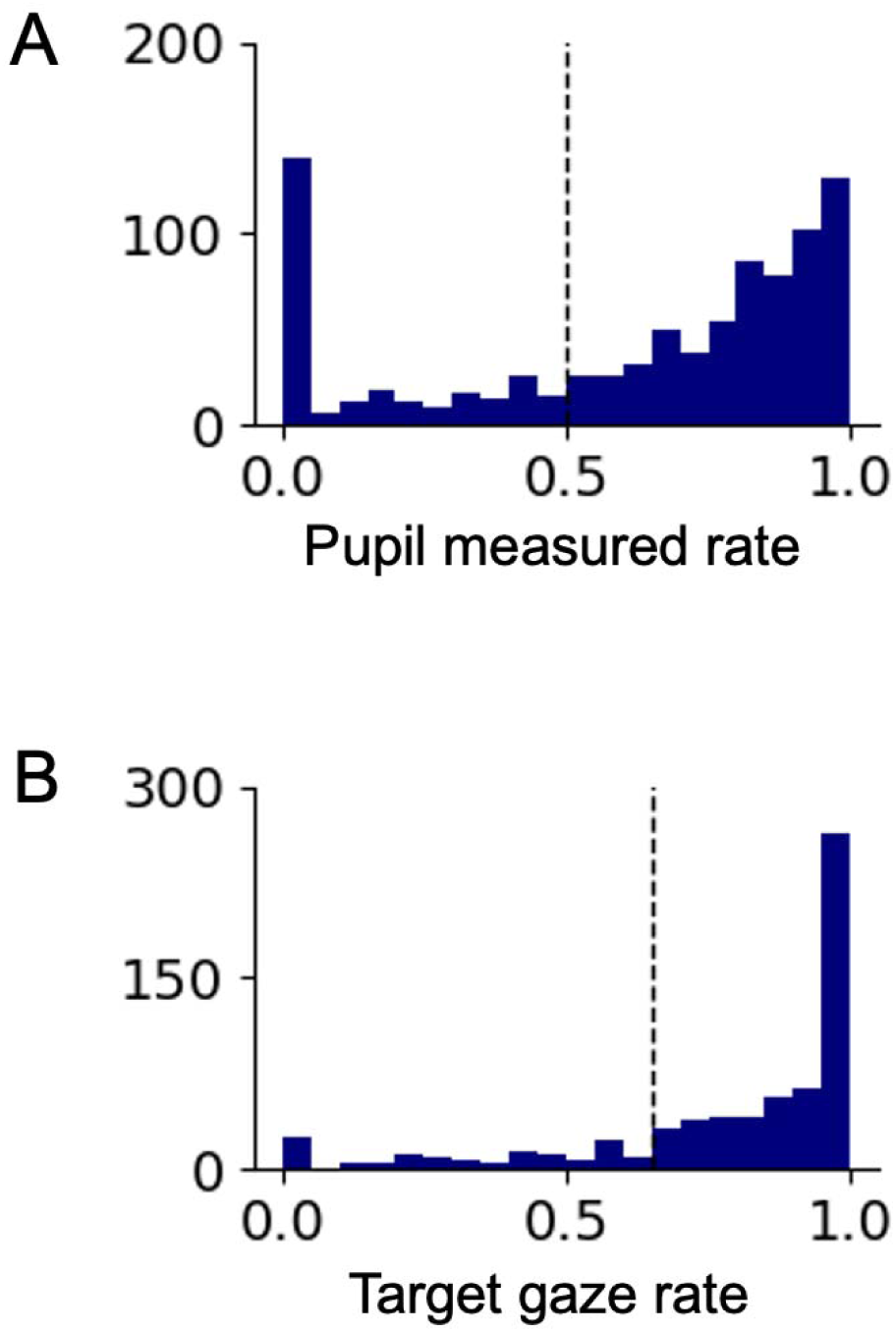
Thresholds for pupil dilation analysis. (**A**) The rate of intervals during which pupil diameter was measured between 2-seconds before and 4-seconds after the selected movement onsets. The dashed line is the threshold for the subsequent pupil dilation analysis. (**B**) The rate of target gaze between 2-seconds before and 4-seconds after the selected movement onsets. The dashed line is the threshold for the subsequent pupil dilation analysis.

**Fig. S10.**
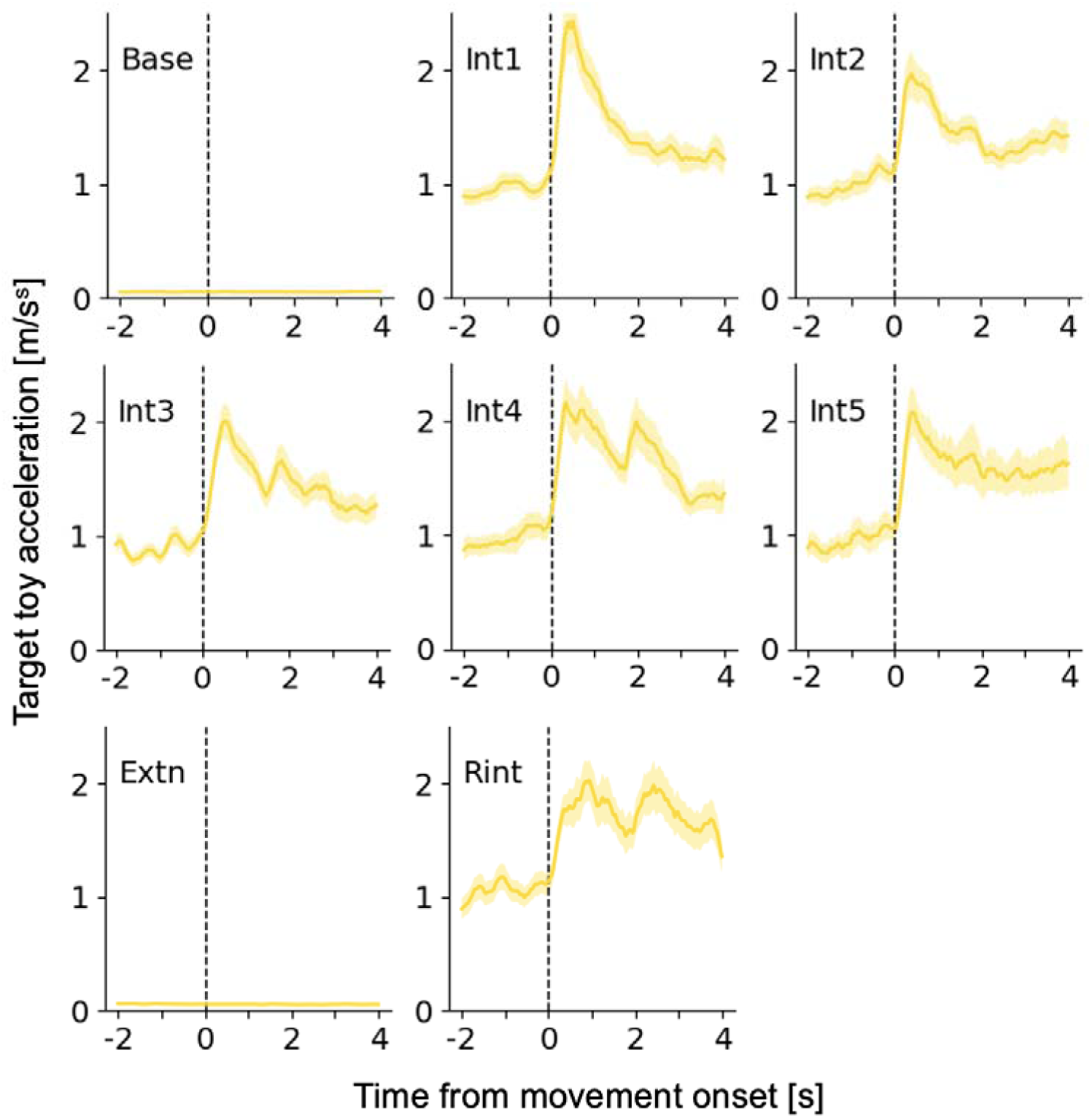
Accelerations of the target toy associated with the onset of arm movements. In the baseline and extinction phases, the target toy was completely stopped. In the interaction and reinteraction phases, a peak of acceleration was observed immediately after the onset of arm movements, while the target toy continued to swing. The used trials were the same as pupil dilation analysis (table. S2).

**Fig. S11.**
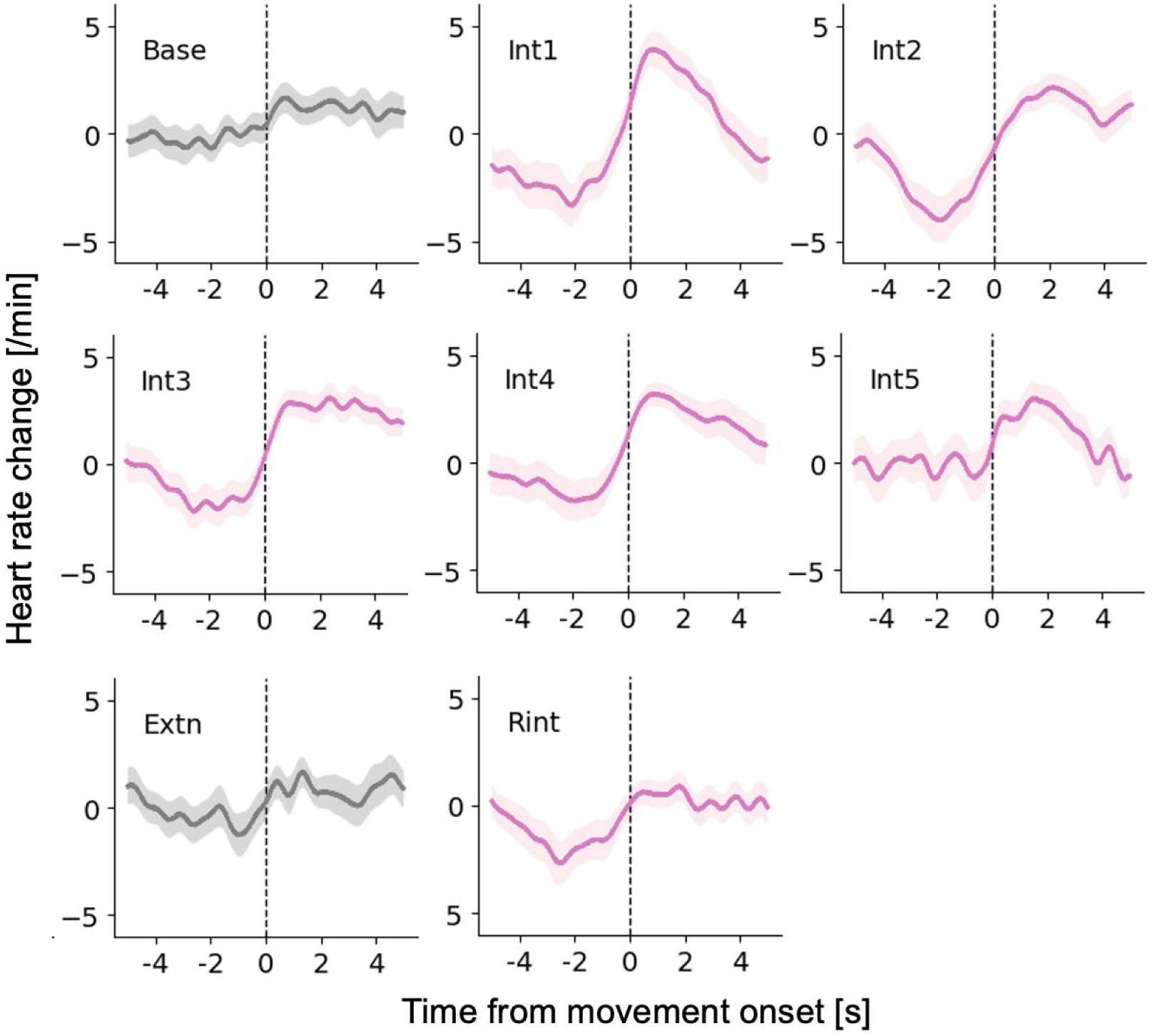
Heart rate changes associated with the onset of arm movements. No clear increase or decrease in heart rate was observed during the baseline phase. However, after the mobile connection, heart rates began to exhibit two characteristic variations. One was a decrease before the onset of movement. the other was an increase at the moment of the movement onset. These changes were prominent in the early part of the interaction phase and gradually diminished in the later part. The used trials were the same as pupil dilation analysis (table. S2). Since heart rate changes more slowly than pupil diameter, it is possible that the observed changes reflect the contamination of the preceding movement.

**Fig. S12.**
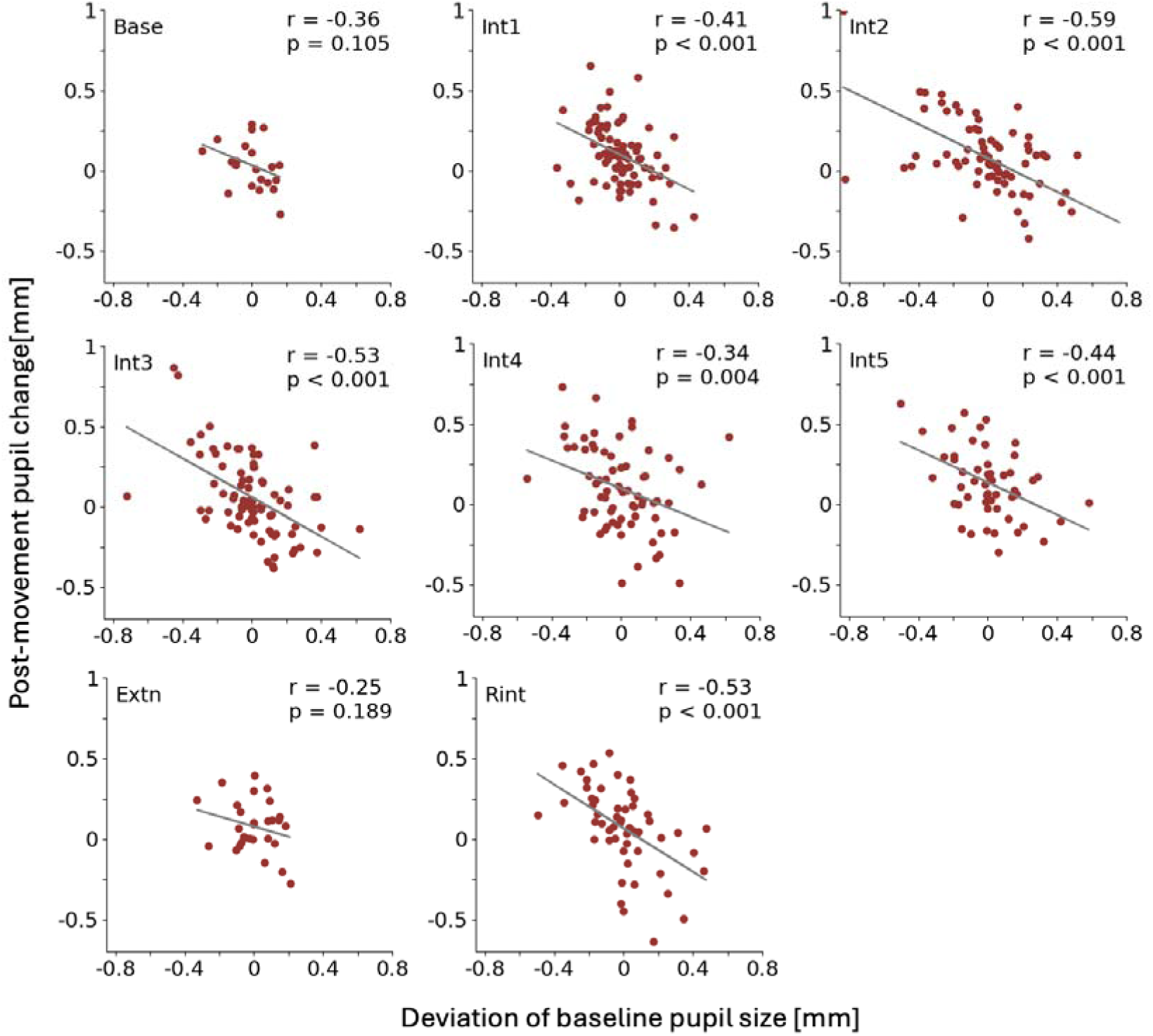
Correlation analysis between baseline pupil size and post-movement change. X-axis represents the deviation of baseline pupil size from each individual average. Baseline pupil size was calculated by averaging pupil diameters in the interval from 2 to 1 seconds before the movement onset. Y-axis represents the degree of change in pupil size averaged in the interval from 1 to 3 seconds after the movement onset. Moderate negative correlations were observed throughout the interaction and reinteraction phases, indicating that the degree of post-movement pupil dilation was smaller when the baseline pupil size was larger. However, no significant correlations were observed in the baseline and extinction phases.

**Fig. S13.**
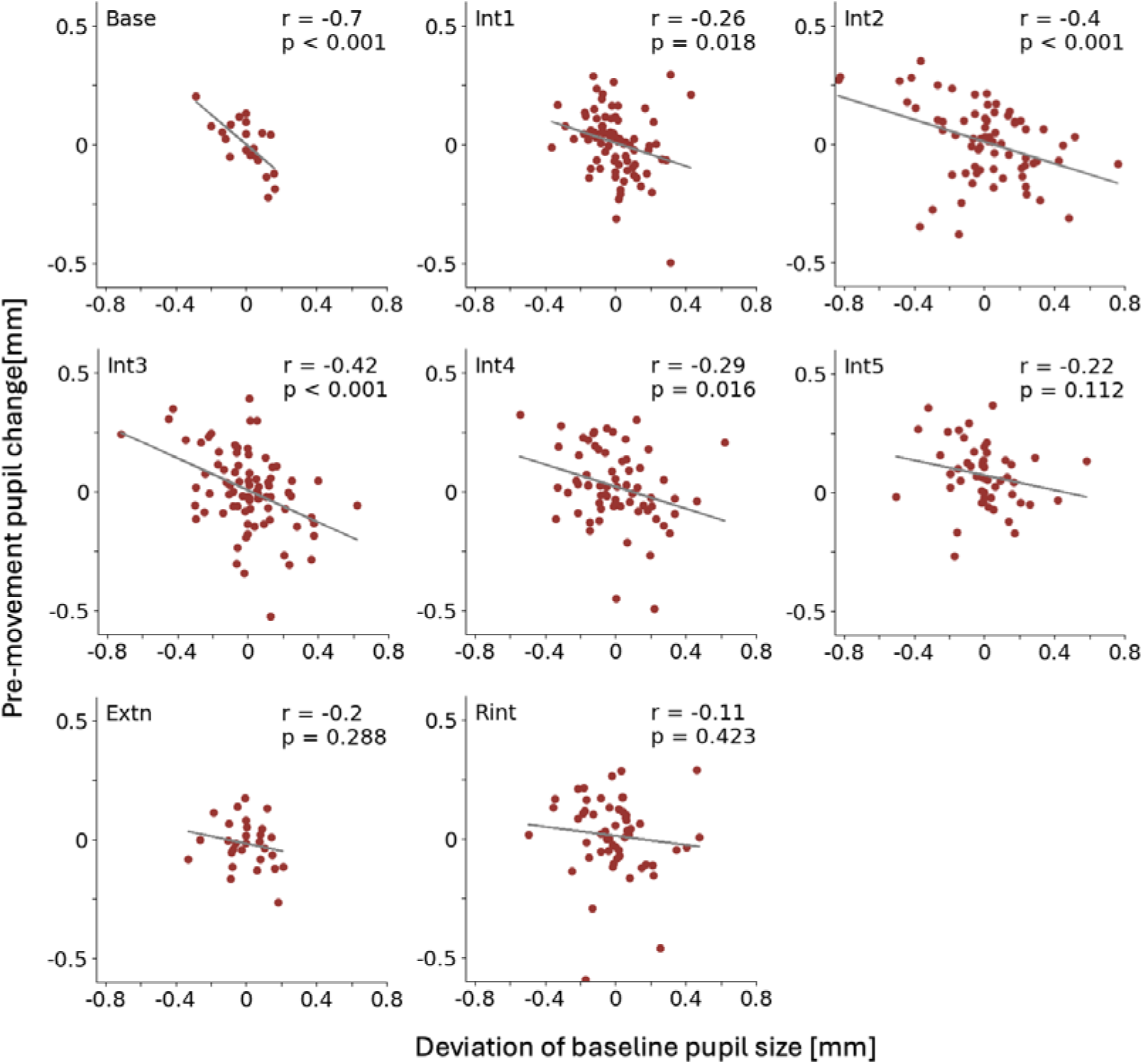
Correlation analysis between baseline pupil size and pre-movement change. X-axis represents the deviation of baseline pupil size from each individual average. Y-axis represents the degree of change in pupil size averaged in the interval from 1 to 0 seconds before the movement onset. A strong negative correlation was observed in the baseline phase, and moderate negative correlations were found in the Int2 and Int3 phases. However, no significant correlations were observed in the later part of the measurement. Notably, the pre-movement pupil dilation in the Int5 phase did not correlate with baseline pupil sizes.

**Fig. S14.**
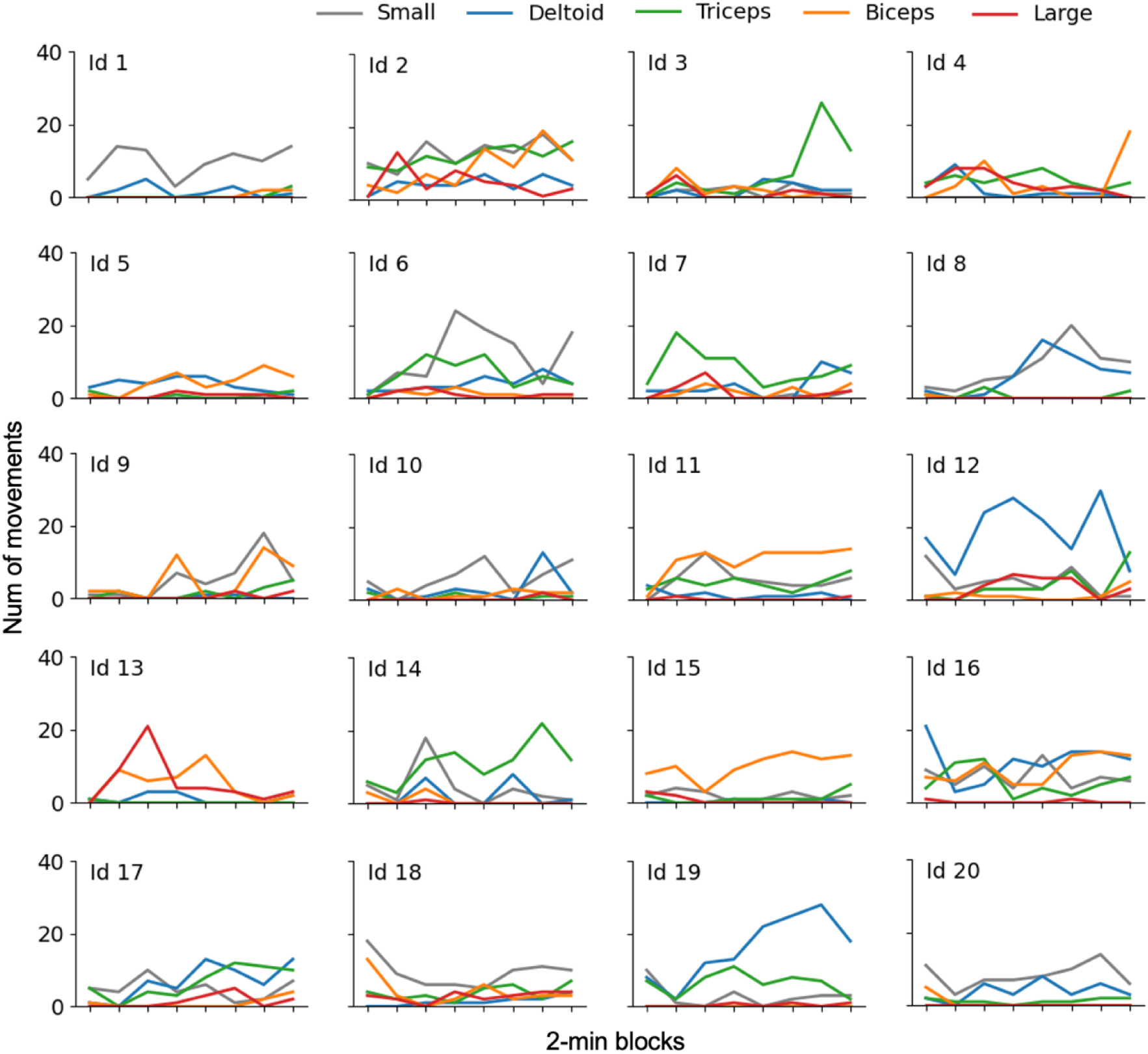
Individual plots for the number of movements in each cluster. There was substantial individual variation in the cluster composition. Some infants (Id 12 and 19) mainly used the deltoid muscle, while others (Id 3 and Id 14) used the triceps. Additionally, some infants (Id 11 and Id 15) also used the biceps.

**Fig. S15.**
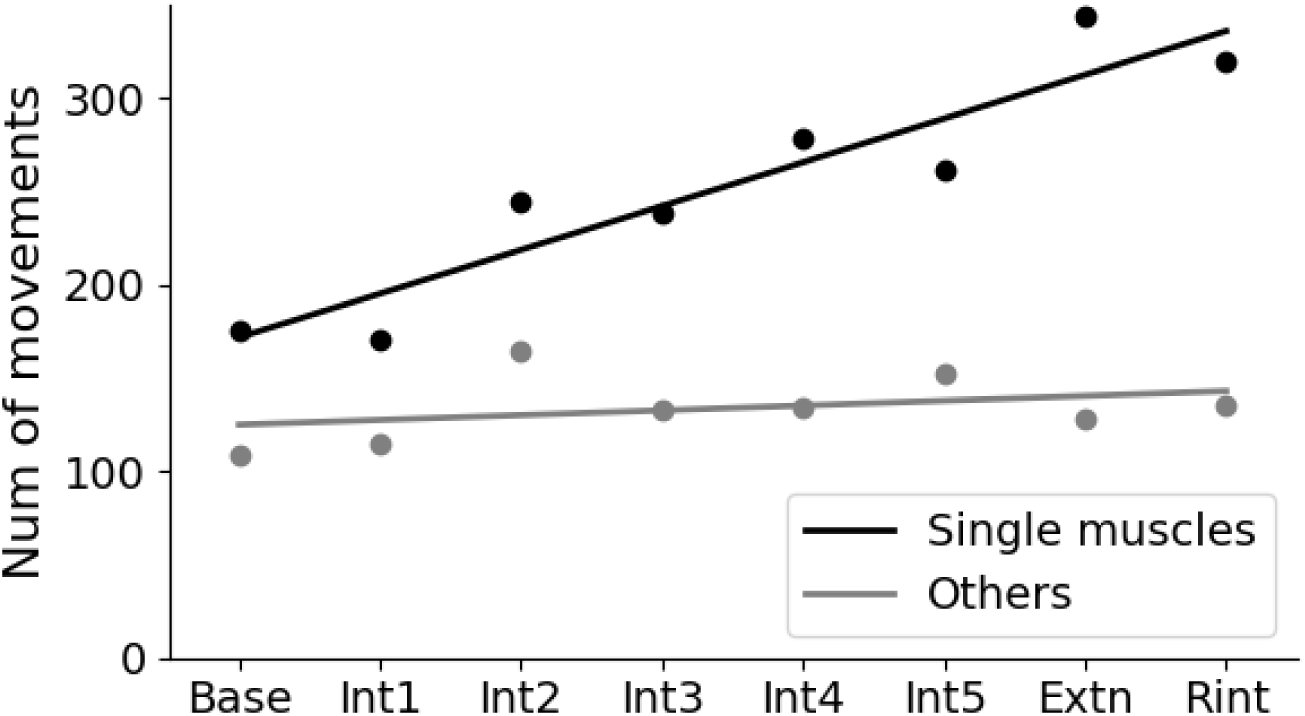
Changes in the number of movements grouped by muscular activity patterns. Single-muscle group includes the movements clustered as deltoid, biceps, and triceps (Fig. 3A-C). The other group includes the movements clustered as small and large. Each dot represents the number of movements summed across infants in each phase, shown together with the regression line. The single-muscle group exhibited a greater increase in movements relative to the other group.

**Table S1.** The number of movements in each infant. ‘Original’ is the number of detected movements using EMG signals. ‘Interval selection’ is the number of remaining trials after excluding those with prior movements within a 6-second window. ‘Pupil selection’ is the number of remaining trials after additional selection based on the measured rate of pupil diameter and the rate of toy gazing.

| ID | Original | Interval selection | Pupil selection |
| --- | --- | --- | --- |
| 1 | 99 | 48 | 22 |
| 2 | 339 | 32 | 17 |
| 3 | 109 | 39 | 13 |
| 4 | 119 | 50 | 21 |
| 5 | 76 | 38 | 13 |
| 6 | 195 | 54 | 23 |
| 7 | 124 | 50 | 30 |
| 8 | 127 | 48 | 2 |
| 9 | 102 | 28 | 18 |
| 10 | 92 | 35 | 14 |
| 11 | 182 | 53 | 41 |
| 12 | 258 | 48 | 32 |
| 13 | 95 | 32 | 15 |
| 14 | 148 | 41 | 15 |
| 15 | 115 | 54 | 28 |
| 16 | 271 | 41 | 18 |
| 17 | 165 | 48 | 28 |
| 18 | 170 | 52 | 41 |
| 19 | 205 | 55 | 46 |
| 20 | 112 | 46 | 27 |
| **Total** | **3103** | **892** | **464** |

**Table S2.** The number of movements used for pupil dilation analysis in each phase.

| ID | Base | Int1 | Int2 | Int3 | Int4 | Int5 | Extn | Rint |
| --- | --- | --- | --- | --- | --- | --- | --- | --- |
| 1 | 3 | 0 | 5 | 2 | 8 | 3 | 0 | 4 |
| 2 | 0 | 5 | 4 | 3 | 1 | 1 | 0 | 0 |
| 3 | 0 | 8 | 3 | 0 | 1 | 0 | 1 | 0 |
| 4 | 0 | 2 | 3 | 8 | 5 | 1 | 0 | 2 |
| 5 | 0 | 2 | 2 | 1 | 0 | 2 | 0 | 6 |
| 6 | 1 | 4 | 2 | 4 | 6 | 4 | 0 | 2 |
| 7 | 0 | 9 | 5 | 10 | 2 | 1 | 0 | 3 |
| 8 | 0 | 1 | 0 | 0 | 0 | 0 | 1 | 0 |
| 9 | 1 | 1 | 0 | 7 | 3 | 1 | 0 | 5 |
| 10 | 0 | 1 | 2 | 4 | 3 | 2 | 0 | 2 |
| 11 | 0 | 7 | 7 | 7 | 6 | 6 | 4 | 4 |
| 12 | 2 | 7 | 4 | 3 | 3 | 7 | 3 | 3 |
| 13 | 0 | 1 | 2 | 7 | 2 | 2 | 0 | 1 |
| 14 | 3 | 2 | 2 | 2 | 2 | 2 | 2 | 0 |
| 15 | 5 | 7 | 5 | 1 | 5 | 1 | 3 | 1 |
| 16 | 0 | 7 | 5 | 3 | 0 | 0 | 3 | 0 |
| 17 | 1 | 3 | 5 | 2 | 5 | 7 | 0 | 5 |
| 18 | 2 | 7 | 2 | 5 | 6 | 5 | 7 | 7 |
| 19 | 2 | 4 | 9 | 10 | 7 | 4 | 2 | 8 |
| 20 | 1 | 3 | 5 | 3 | 4 | 5 | 3 | 3 |
| **Total** | 21 | 81 | 72 | 82 | 69 | 54 | 29 | 56 |

**Table S3.** The number of movements divided by clusters used for pupil dilation analysis. Single-muscle group includes the movements clustered as deltoid, biceps, and triceps (Fig. 3A-C). The other group includes the movements clustered as small and large.

| ID | Int4<br>Single<br>muscles | Int4<br>Others | Int5<br>Single<br>muscles | Int5<br>Others |
| --- | --- | --- | --- | --- |
| 1 | 2 | 6 | 1 | 2 |
| 2 | 0 | 1 | 0 | 1 |
| 3 | 0 | 1 | 0 | 0 |
| 4 | 4 | 1 | 0 | 1 |
| 5 | 0 | 0 | 2 | 0 |
| 6 | 4 | 2 | 4 | 0 |
| 7 | 1 | 1 | 0 | 1 |
| 8 | 0 | 0 | 0 | 0 |
| 9 | 2 | 1 | 1 | 0 |
| 10 | 3 | 0 | 1 | 1 |
| 11 | 4 | 2 | 4 | 2 |
| 12 | 1 | 2 | 7 | 0 |
| 13 | 2 | 0 | 2 | 0 |
| 14 | 2 | 0 | 2 | 0 |
| 15 | 4 | 1 | 1 | 0 |
| 16 | 0 | 0 | 0 | 0 |
| 17 | 4 | 0 | 6 | 1 |
| 18 | 4 | 2 | 2 | 3 |
| 19 | 2 | 5 | 3 | 1 |
| 20 | 2 | 2 | 2 | 3 |
| Total | 41 | 27 | 38 | 16 |

